# A structure-informed evolutionary model for predicting viral immune escape and evolution

**DOI:** 10.1101/2025.07.31.667864

**Authors:** Chonghao Wang, Lu Zhang

## Abstract

Persistent emergence of viral variants capable of evading host immunity constitutes a significant threat to public health. This antigenic evolution frequently outpaces the development of vaccines and therapeutics, highlighting the necessity of predictive models surveilling the immune escape potential of emerging variants. However, existing models suffer from two key limitations: they inadequately incorporate protein structural information and neglect the importance of distinguishing large-impact mutations from neutral ones given multiple mutations. To address these gaps, we presented KEScape, a deep learning model designed to predict viral immune escape and evolution. KEScape integrates evolutionary context with protein structural information, introduces a novel top-K ***L***_**2**_-differential pooling mechanism to prioritize mutations with large functional effects, and incorporates a supervised ***L***_**2**_ margin loss to facilitate the ***L***_**2**_-distance-based ranking of high-impact mutations. We demonstrated that KEScape significantly outperformed state-of-the-art models on the most comprehensive benchmark to date, comprising eleven deep mutational scanning experiments spanning diverse viruses. Furthermore, KEScape exhibited outstanding performance in the practical applications of identifying immune escape hotspots and variants in emerging lineages and real-time surveillance of lineages associated with WHO-designated variants of SARS-CoV-2. These results show that KEScape is an effective model to predict viral immune escape and evolution. Its capacity for early warning can directly inform public health interventions and guide the development of countermeasures, thereby mitigating the threat of future viral pandemics.

## Introduction

The co-evolutionary history of humans and viruses is marked by recurrent pandemics that have inflicted catastrophic mortality and shaped societal development [1]. For instance, the 1918 influenza pandemic, caused by the H1N1 influenza A virus, resulted in an estimated 50 million deaths worldwide [2]. More recently, severe acute respiratory syndrome coronavirus 2 (SARS-CoV-2) emerged as a global health crisis, precipitating a pandemic that has claimed millions of lives [3, 4]. These historical precedents establish a clear and urgent mandate: to develop strategies that can mitigate the impact of future viral threats. To this end, the World Health Organization (WHO) introduced the concept of “Disease X” – a framework conceptualizes the challenge of preparing for a pandemic caused by a yet-to-be-identified pathogen. This proactive stance emphasizes global surveillance, which focuses on threats with high pandemic potential, such as highly pathogenic avian influenza (H5N1) and other zoonotic agents that have demonstrated the capacity to cross the species barrier [5–7]. Given the broad consensus that future pandemics are inevitable, pandemic preparedness must shift from a primarily reactive paradigm to a proactive one. A robust preparedness framework should not only integrate systematic pathogen genome sequencing and rapid vaccine development, but also include a critical third pillar: the timely identification of emerging viral variants with immune escape potential in the genomic surveillance. The capacity to recognize such variants at an early stage is essential for prioritizing experimental characterization, guiding vaccine and therapeutic updates, and enabling immediate public health interventions [8–13].

The persistence of viral threats is fundamentally shaped by the complex coevolutionary dynamics between viruses and their hosts. During the initial emergence of a novel virus in an immunologically naive population, viral spread is primarily driven by factors such as intrinsic transmissibility, host susceptibility, and ecological or epidemiological conditions. Once substantial population-level immunity has been established, for example through a large wave of infection, widespread vaccination, or both, the evolutionary landscape shifts. This newly established population-level immunity imposes intense selective pressure on the viral population. In turn, viruses continuously evolve, accumulating mutations that facilitate evasion of host immune surveillance – a phenomenon known as viral immune escape [14–20]. While advancements in immunology have yielded highly effective vaccines that bolster host immunity and save countless lives [21–24], the adaptive antigenic evolution of viruses poses an ongoing challenge. Specifically, viral mutations, particularly in key antigenic sites, can reduce or abrogate the binding affinity of vaccine-induced antibodies, thereby diminishing vaccine efficacy. This necessitates periodic reformulation of vaccines to match circulating viral lineages. It poses a critical challenge to accurately predict which viral variants will escape existing population immunity and achieve future prevalence. Current practices, such as the WHO’s influenza vaccine composition meetings, rely on expert consultation. However, mismatches between vaccine lineages and dominant circulating lineages have occurred, leading to suboptimal vaccine effectiveness in several seasons [25–27].

To better understand the landscape of potential escape mutations, experimental techniques like deep mutational scanning (DMS) have been developed [28]. DMS 2 allows for the high-throughput functional characterization of numerous viral mutants, assessing their impact on properties such as antibody binding or viral fitness. However, DMS experiments are resource-intensive and inherently limited in scope relative to the vast combinatorial space of possible mutations. For example, for a protein of length *n*, there are 19 × *n* possible single amino acid substitutions, and exponentially more multiple mutations. Computational approaches therefore provide a promising means of complementing experimental characterization. For example, DMS experiments performed during an initial wave of infection or following early vaccination campaigns can quantify the effects of mutations relative to the wild-type virus. These data can then be used to train computational models capable of predicting the immune escape potential of subsequently emerging variants.

Several computational models have been developed to predict viral immune escape. For instance, Hie et al. [29] introduced CSCS, a model to predict viral escape by analyzing the differences between the wild-type and mutant sequences in the embedding spaces generated by a protein language model. The core premise of CSCS is that viable escape variants must maintain essential fitness while exhibiting sufficient divergence from the wild-type virus to evade immune surveillance. MLAEP [30] leverages a genetic algorithm to identify potential immune escape variants of SARS-CoV-2 by simulating natural selection. The genetic search is informed by a fitness landscape generated by integrating sequence features with structural information. The sequence feature is derived from the receptor-binding domain (RBD), which is encoded using ESM-2, a protein language model (PLM). The structural information, pertaining to ACE2 and antibody complexes, is processed by a structure-aware transformer. Another prominent model, EVEscape [31], operates on the assumptions that an escape variant should preserve its fitness, while simultaneously disrupting the antibody binding interface on the viral protein surface. To quantify escape potential, EVEscape integrates three distinct metrics: fitness, accessibility, and dissimilarity. E2VD [32] utilizes a pre-trained PLM to extract residue-level embeddings, and employs a convolution neural network and an attention block to capture the local and global dependencies of residues. E2VD proposes a multi-task framework to address the challenge of imbalanced data between beneficial and deleterious mutations, which consists of three tasks: receptor-binding, expression, and immune escape.

While these models have advanced the field of viral immune escape prediction, they either omit protein structural information or derive a limited set of static features from protein structures. However, structural changes induced by mutations are demonstrated to be a primary mechanism of viral immune escape [33, 34]. In addition, most models for predicting mutational effects overlook the importance of distinguishing the signals of large-effect mutations from the noise introduced by neutral mutations in variants carrying multiple mutations. Given that viral fitness and immune escape are driven by a small number of mutations with large functional effects [19], while the majority are phenotypically neutral or attenuated [35–37], the noise from abundant neutral mutations can obscure the signals from determinative ones.

In response to these gaps, we propose KEScape, a model to predict viral immune escape and evolution. The primary application of KEScape is to predict the immune escape potential of emerging variants when population immunity constitutes a major selective pressure and DMS datasets from the same protein family as the target viral protein are available. To make predictions, KEScape first integrates evolutionary and structural information into unified embeddings, and then models residue-residue dependencies to generate context-aware unified embeddings. Subsequently, the embedding pairs corresponding to the *K* mutations with the largest functional impacts, as quantified by their *L*_2_ distances, are selectively pooled and used for prediction (Fig. 1A). KEScape introduces three innovative features: (i) a fusion mechanism that integrates evolutionary embeddings and structural embeddings into unified embeddings; (ii) a top-K *L*_2_-differential pooling mechanism to prioritize mutations with significant functional impacts; (iii) a supervised *L*_2_ margin loss that encourages the model to enlarge *L*_2_ distances between immune escape variants and their corresponding wild-type references in the embedding space.

**Fig. 1:**
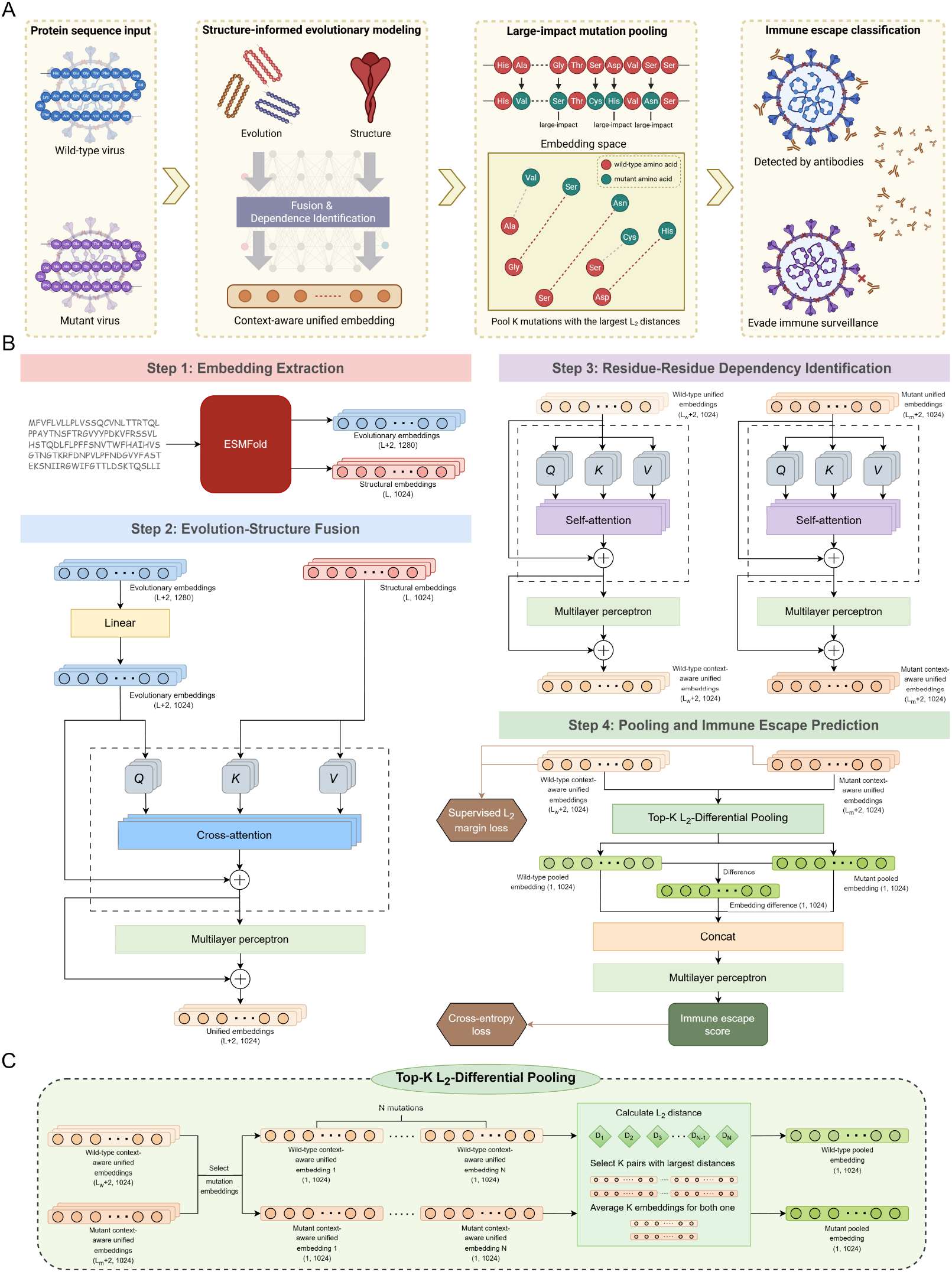
Overview of KEScape. (**A**) An illustrative diagram of KEScape. The model processes protein sequences of the wild-type and mutant viruses as inputs, generating context-aware unified embeddings by integrating evolutionary and structural embeddings to capture residue-residue dependencies. For each sequence, KEScape selects and averages the pairs of context-aware unified embeddings (*K* = 5) corresponding to *K* mutations with the largest *L*_2_ distances. Subsequently, KEScape estimates the likelihood of the mutant virus evading host immune surveillance. (**B**) KEScape architecture. KEScape (i) derives evolutionary embeddings and structural embeddings from ESMFold for an input protein sequence; (ii) fuses these embeddings into unified embeddings via a cross-attention module with residual connections; (iii) captures residue-residue dependencies through a self-attention module; (iv) employs top-K *L*_2_-differential pooling to focus on mutations with the largest functional impacts and predicts an immune escape score for the mutant sequence; (v) calculates the supervised *L*_2_ margin loss and weighted cross-entropy loss to evaluate the predictive performance during the training stage. (**C**) Top-K *L*_2_-differential pooling mechanism. Given *N* mutations, the pooling mechanism calculates *L*_2_ distances for *N* pairs of wild-type and mutant context-aware unified embeddings, selects *K* pairs with largest *L*_2_ distances, and averages the corresponding *K* wild-type and mutant embeddings, respectively.

To address the limited scope of prior benchmarks, which have typically been confined to five or fewer viral species, we evaluated KEScape using a comprehensive suite of eleven DMS datasets. To our knowledge, this benchmark represents the most extensive collection assembled to date, spanning a diverse range of viruses and effectively doubling the scale of prior evaluations. In this benchmark, KEScape outperformed all competing models, achieving a mean area under the precision-recall curve (AUPRC) of 0.534 compared to 0.365 from the second best-performing model. Extensive studies have demonstrated that viruses frequently acquire mutations within specific regions and sites [15, 38, 39] to evade host immune surveillance. Consequently, identifying such mutation hotspots in the emerging lineages provides a valuable probe to assess their immune escape potential during pandemics. Therefore, we examined the performance of KEScape in identifying immune escape hotspots on the SARS-CoV-2 BA.2 spike protein and the results were consistent with previous independent studies [20, 40–42]. In addition, we demonstrated that, when trained on SARS-CoV-2 Wuhan-Hu-1 and other viral proteins, KEScape can identify immune escape variants of SARS-CoV-2 XBB.1.5 lineages. However, we note that KEScape is not recommended for application to viral proteins whose corresponding protein families are not represented in the training set, as predictions in such settings may be unreliable. During epidemics and pandemics, novel viral variants frequently emerge across distinct geographic regions. To prepare for subsequent waves of re-infection, accurately predicting the immune escape potential of emerging variants in a large scale is essential. We evaluated KEScape using temporally partitioned sequences obtained from the GISAID database [31, 43]. The assessment demonstrated that KEScape could identify lineages associated with WHO-designated variants of SARS-CoV-2 (hereafter referred to as WDV lineages) by assigning higher immune escape scores to them compared to contemporaneous non-WDV lineages. To elucidate the internal mechanisms of KEScape, we analyzed its latent embeddings and attention weights, confirming the KEScape’s interpretability. Subsequently, we demonstrated the contribution of innovative components through ablation studies.

## Results

### Overview of KEScape

KEScape comprises four main steps to predict viral immune escape and evolution. First, it employs ESMFold [44] to generate evolutionary and structural embeddings for both the wild-type and mutant viral protein sequences (Fig. 1B step 1). In step 2, KEScape leverages a multi-head cross-attention module to systematically integrate structural and evolutionary embeddings. Within this framework, evolutionary embeddings serve as queries, while structural embeddings are keys and values, selectively amplified and combined according to their relevance in modeling immune escape for each query (Methods). After residual connections, KEScape generates unified embeddings by incorporating these reweighted structural embeddings into evolutionary embeddings (Fig. 1B step 2). The fusion is applied to both the wild-type and mutant sequences. In step 3 (Fig. 1B step 3), KEScape generates context-aware unified embeddings by capturing residue-residue dependencies from unified embeddings based on a self-attention module, enabling the modeling of intra-sequence interactions. In step 4, the context-aware unified embeddings of the wild-type and mutant sequences are put into the top-K *L*_2_-differential pooling module (Fig. 1C and supplementary notes). Given *N* mutations, this module first isolates *N* pairs of context-aware unified embeddings corresponding to the mutated residues in both sequences. Pairwise *L*_2_ distances are computed between the *N* wild-type/mutant embedding pairs, after which the *K* pairs exhibiting maximal divergence (*K* empirically determined as min(5, *N*)) are retained. The *K* retained embedding pairs are averaged, generating two pooled embeddings encoding the mutational effects from these *K* mutations for the wild-type and mutant sequences, respectively. These two pooled embeddings and their differences are concatenated and processed through a multilayer perceptron to predict an immune escape score for the mutant sequence (Fig. 1B step 4). The loss function of KEScape comprises two components: (i) a supervised *L*_2_ margin loss, which increases the *L*_2_ distance between the wild-type and mutant embeddings of escape variants while reducing the corresponding distance for non-escape variants; (ii) a weighted cross-entropy loss, which evaluates the classification performance.

### KEScape demonstrates state-of-the-art concordance with DMS experiments

We collected eleven DMS datasets spanning diverse viral proteins [31, 45–52] (Supplementary Table 1) and benchmarked the performance of KEScape against five computational models: CSCS, EVEscape, fine-tuned CSCS, fine-tuned EVEscape, and MLAEP. Fine-tuned versions of CSCS and EVEscape were included to control for the performance gains attributable to the training datasets, as both CSCS and EVEscape are originally unsupervised models. Because MLAEP is specifically designed for SARS-CoV-2, its performance was evaluated only on SARS-CoV-2 DMS datasets. In addition, the benchmarking against E2VD was conducted using its own designated dataset, as the antibody sequences it required were unavailable in our collected datasets.

Comparative analysis across eleven DMS datasets (Fig. 2A, C) revealed the superior performance of KEScape, which achieved significantly higher area under the receiver operating characteristic curve (AUROC) than other models. Specifically, KEScape attained a mean AUROC of 0.901, compared with 0.851 for fine-tuned CSCS, 0.751 for fine-tuned EVEscape, 0.682 for EVEscape, and 0.587 for CSCS. As all datasets were imbalanced, we also evaluated model performance using AUPRC. KEScape likewise significantly outperformed the competing models in terms of AUPRC, achieving a mean AUPRC of 0.534, whereas fine-tuned CSCS, fine-tuned EVEscape, EVEscape, and CSCS achieved mean AUPRC values of 0.365, 0.168, 0.125, and 0.064, respectively (Fig. 2A, D).

**Fig. 2:**
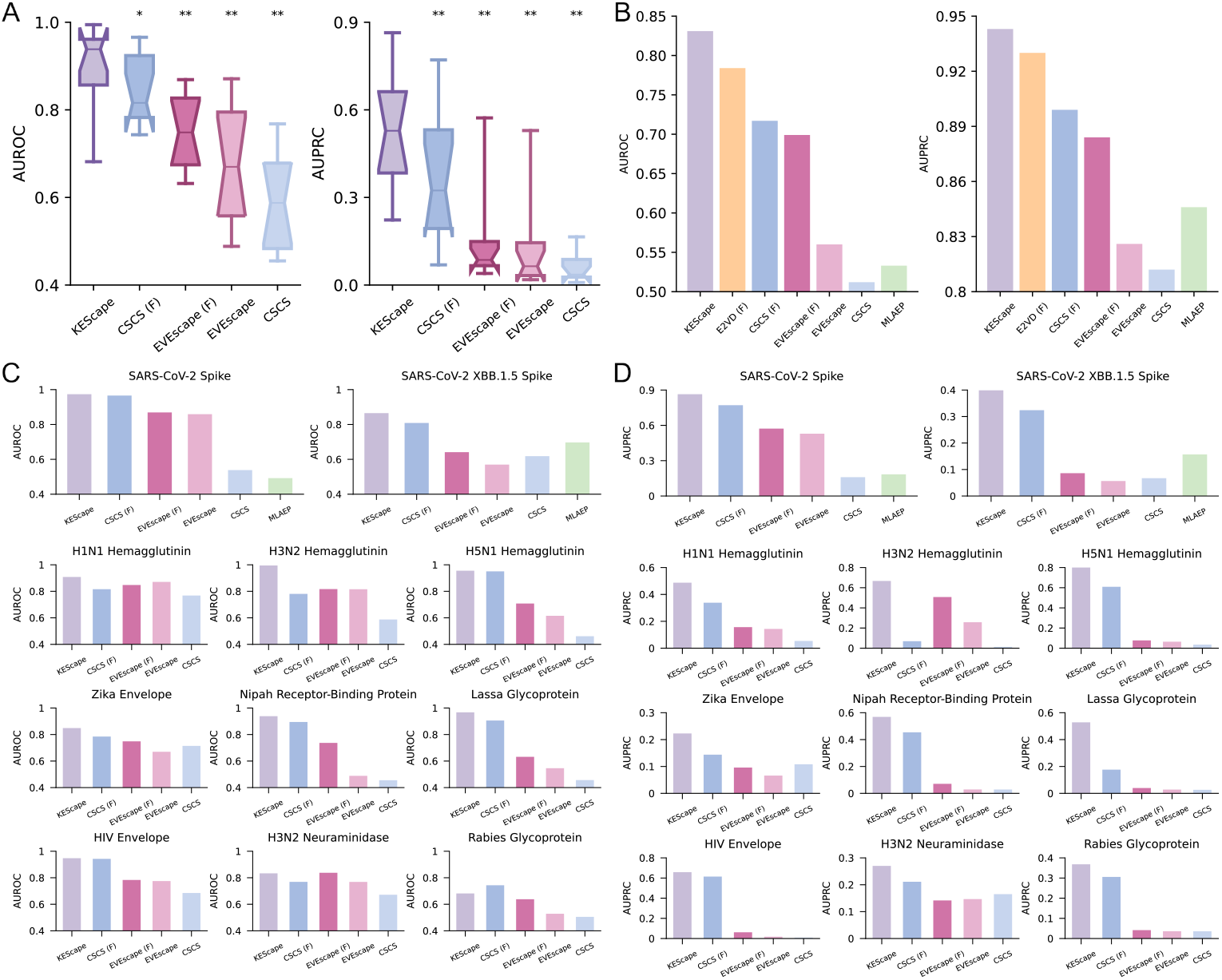
Comparative performance evaluation of KEScape across diverse DMS datasets. (**A**) Aggregate comparison of AUROC and AUPRC values between KEScape and state-of-the-art computational models for viral immune escape prediction across eleven DMS datasets. The suffix ‘(F)’ designates the fine-tuned version of a respective model (e.g., CSCS (F)). Symbols above the competing models indicate the statistical significance of the extent to which KEScape outperforms each corresponding model: *n*.*s*. denotes *p* > 0.05; ∗ denotes 0.01 < *p* ≤ 0.05; and ∗∗ denotes *p* ≤ 0.01. The *p*-values were obtained using paired *t*-tests. (**B**) Extended benchmark including E2VD, assessed on its SARS-CoV-2 DMS dataset containing antibody sequences. E2VD requires both viral protein sequences and antibody sequences as inputs, whereas other models operate solely on viral protein inputs. (**C**) AUROC comparison of KEScape and competing state-of-the-art models across the eleven DMS datasets. MLAEP was evaluated only on the SARS-CoV-2 spike and SARS-CoV-2 XBB.1.5 spike datasets because it was specifically designed for SARS-CoV-2. (**D**) AUPRC comparison of KEScape and competing state-of-the-art models across the eleven DMS datasets.

Notably, we found that all evaluated models exhibited variable performance across different datasets (Fig. 2A, C, D). KEScape achieved maximal AUPRC (0.864) for SARS-CoV-2 spike, while showed reduced AUPRC (0.223) for Zika envelope. Similar trends were observed for other models. For example, fine-tuned CSCS and fine-tuned EVEscape, which both reached their highest AUPRC on SARS-CoV-2 spike (0.771 and 0.572, respectively) but underperformed on H3N2 hemagglutinin and Lassa glycoprotein. This heterogeneity indicates that different viral proteins exhibit distinct propensities for specific immune escape mechanisms, which are not fully captured by current computational models. For instance, glycan shielding was reported critical for HIV immune escape [31]. Furthermore, we observed consistent performance patterns across models for certain datasets, with uniformly strong or weak predictive accuracy observed in specific viral protein contexts. For example, KEScape, fine-tuned CSCS, fine-tuned EVEscape and EVEscape all excelled in SARS-CoV-2 spike dataset, while exhibiting comparatively reduced performance in the rabies glycoprotein dataset. In summary, these results suggest that the difficulty of predicting immune escape varies substantially across different viral proteins. Beyond potential data noise, the variation likely arises from differences in immune escape mechanisms, with certain viral proteins employing more complex mechanisms.

To benchmark KEScape against E2VD, we used a SARS-CoV-2 DMS dataset containing antibody sequences provided by the E2VD study [32]. As illustrated in Fig. 2B, KEScape outperformed all competing models, including E2VD. We note that while E2VD explicitly requires antibody sequence information as input, KEScape and other models operate without this input. Our results reveal that KEScape can achieve superior performance in the absence of antibody-specific information, implying that its structure-informed evolutionary modeling - operating solely through viral protein analysis - is sufficient to capture critical determinants of viral immune escape.

### KEScape identifies immune escape hotspots and variants in emerging lineages

A practical assessment of model performance is whether it can identify immune escape hotspots and variants in emerging lineages that were not represented in the training set. To evaluate hotspot identification, we leveraged the well-characterized evolutionary trajectory of SARS-CoV-2 and selected the BA.2 lineage, which was not in the training set, as a test case for assessing KEScape’s ability to identify immune escape hotspots. Mapping the max per-residue KEScape scores (representing the highest KEScape score among all single amino acid substitutions in each residue, see Methods) onto the SARS-CoV-2 BA.2 spike protein structure (Protein Data Bank (PDB): 7XIX), this revealed that residues with high scores predominantly clustered in RBD and the N-terminal domain (NTD), while most of residues outside these two domains exhibited mild predicted impact on immune escape (Fig. 3A, B, Supplementary Fig. 1). These findings align with the established evidence that RBD and NTD mutations are primary drivers of immune escape in Omicron variants [20, 53, 54]. Notably, many residues with high KEScape scores correspond to experimentally validated immune escape hotspots. For instance, residue 483 (align with residue 486 in SARS-CoV-2 Wuhan-Hu-1) is a well-documented site critical for immune escape [40–42]. This residue constitutes a primary target for several neutralizing antibodies [55, 56], and the F486V mutation present in BA.4/5 facilitates escape from multiple antibodies [20, 40]. Similarly, mutations in residue 343 (346 in Wuhan-Hu-1), present in several Omicron sublineages, confer significant antigenic variation [53]. Furthermore, other high-scoring residues—including 142 (145), 441 (444), 449 (452), 453 (456), and 457 (460)—reside within functionally critical regions and can be associated with viral immune escape [20, 57–61].

**Fig. 3:**
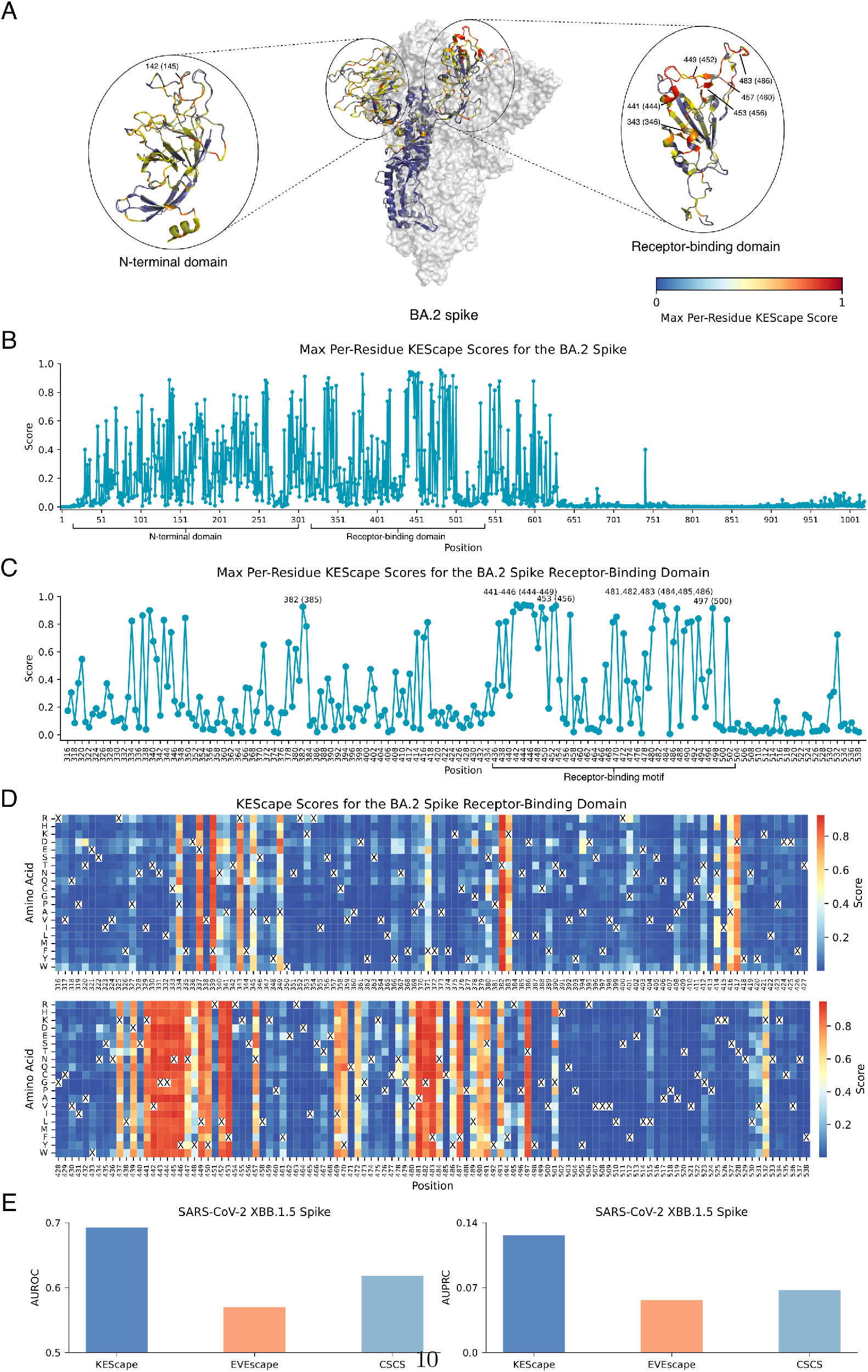
KEScape identifies immune escape hotspots and variants in spike proteins from SARS-CoV-2 lineages not represented in the training set. (**A**) Max per-residue KEScape scores mapped onto one chain of the BA.2 spike (pdb: 7XIX). Higher scores, indicated by warmer colors, signify greater potential for immune escape. A subset of residues with high scores is annotated, with their corresponding Wuhan-Hu-1 coordinates indicated in parentheses. (**B**) Max per-residue KEScape scores for the BA.2 spike. (**C**) Max per-residue KEScape scores for the RBD of the BA.2 spike. Residues with higher scores are highlighted, with the Wuhan-Hu-1 reference positions provided in parentheses. (**D**) A comprehensive mutation landscape of the BA.2 RBD. The heatmap displays the KEScape score for every possible single amino acid substitution at each residue position (x-axis). The wild-type amino acid at each position is omitted for clarity, while all alternative amino acids are shown with their respective KEScape scores. (**E**) Performance of KEScape, EVEscape, and CSCS on the SARS-CoV-2 XBB.1.5 spike DMS dataset. None of these models was trained on this lineage.

Visualization of max per-residue KEScape scores for the BA.2 spike RBD (Fig. 3C) demonstrated that the highest-scoring residues were largely concentrated in the receptor-binding motif (RBM). RBM, a critical subdomain of the spike protein, mediates direct interaction with the host cell receptor ACE2 [62, 63] and represents a major target for neutralizing antibodies [64, 65]. This observation aligns with experimental evidence highlighting RBM mutations—such as N460K and F486V—as significant contributors to immune evasion [20, 42, 66]. Taken together, these results demonstrate that KEScape effectively identifies key regions governing immune escape and accurately pinpoints established immune escape hotspots. Analysis of the mean per-residue KEScape scores for the BA.2 RBD (Supplementary Fig. 2) revealed a distribution congruent with the max scores, exhibiting similar patterns of regions with large and small scores. However, several residues exhibited high maximum scores coupled with low mean scores. For example, residue 394 (397) displayed a significant discrepancy between its max and mean scores. Further investigation indicated that this residue possessed approximately three mutant amino acids with high immune escape scores, while the remaining substitutions had low scores (Fig. 3D). This observation suggests that while some residues exhibit broad mutational landscapes conducive to immune escape with minimal fitness cost, others are subject to significant constraint, tolerating only a limited subset of mutations, aside from some neutral mutations, without incurring substantial fitness penalties.

In addition to identifying immune escape hotspots, we evaluated the ability of KEScape to detect immune escape variants. Specifically, KEScape was trained on DMS datasets of SARS-CoV-2 spike and other proteins, and its performance was then evaluated on the DMS dataset of the SARS-CoV-2 XBB.1.5 spike protein, a closely related homolog to the training set. We benchmarked KEScape against EVEscape and CSCS and found that KEScape achieved superior performance (Fig. 3E), demonstrating its effectiveness in detecting immune escape variants.

However, we observed that KEScape’s predictive power is contingent upon the presence of homologous protein families within the training set. While KEScape generalized across distantly related homologs of different influenza A hemagglutinin subtypes, its performance deteriorated significantly when applied to the rabies glycoprotein (Supplementary Fig. 3). As a target lacking homologous representation in the training set, this evaluation yielded an AUROC only slightly above 0.5. These results suggest that KEScape should be applied to viral proteins for which proteins from the same family and viral species are represented in the training set. One practical use case is to train the model on proteins from the original lineage during a pandemic and then predict the immune escape potential of subsequently emerging lineages.

### KEScape as a real-time surveillance tool for emerging lineages

A major challenge in predicting the immune escape potential of emerging SARS-CoV-2 lineages is the selection of an appropriate wild-type reference lineage, particularly in the context of heterogeneous population immunity shaped by diverse vaccination histories and prior infections with distinct viral lineages. To address this challenge, we comprehensively evaluated the ability of KEScape to surveil emerging SARS-CoV-2 lineages across seven 90-day intervals encompassing WDV lineages. We used both the ancestral Wuhan-Hu-1 isolate and the earliest WDV lineage from the preceding interval as wild-type references (Methods). All SARS-CoV-2 spike protein sequences were obtained from GISAID [31]. We benchmarked the performance of KEScape against MLAEP. Regarding the other baseline models, neither CSCS nor E2VD can process mutations involving insertions or deletions (indels). Because the dataset for this experiment contains indels, simply ignoring them is methodologically untenable; omitting indels would erroneously collapse distinct variants, including WDV lineages, into identical sequences. Similarly, EVEscape does not directly support indels. Although its underlying EVE model can theoretically be substituted with TranceptEVE [67] to accommodate such mutations, the scripts for calculating immune escape scores for indels have not been made publicly available. In addition, the Trancept model in TranceptEVE was pre-trained on UniRef100, which could contain all SARS-CoV-2 spike variant sequences instead of just one representative sequence in UniRef50. Including it would therefore introduce an unfair comparison.

As shown in Fig. 4A, KEScape evaluated the immune escape potential of each viral variant relative to Wuhan-Hu-1 by comparing the cumulative distribution function of non-WDV lineages with the maximum KEScape scores of contemporaneous WDV lineages. Across the seven intervals, KEScape identified ten WDV lineages with scores exceeding those of 90% of non-WDV lineages, whereas four WDV lineages received scores below this threshold (Fig. 4E). By comparison, MLAEP assigned five WDV lineages scores substantially below the 90% threshold (Fig. 4B, F), indicating significantly weaker performance than KEScape. In addition, we noticed that, beginning in the fourth interval (September 2021 onward), while most WDV lineages received high scores, a large proportion of non-WDV lineages attained comparably high scores for both KEScape and MLAEP. We posited that this phenomenon arose because non-WDV lineages progressively accumulated escape-conferring mutations acquired from prior dominant lineages. Given the fixed wild-type reference (Wuhan-Hu-1), these mutations rendered non-WDV lineages antigenically distinct from the reference. Supporting this hypothesis, Supplementary Fig. 4A demonstrated a marked increase in the proportion of verified immune escape mutations [68] within non-WDV lineages starting in the fourth interval, correlating with the observed rise in high-scoring non-WDV lineages (Fig. 4A-B). Analysis of the immune escape mutations in WDV lineages (Supplementary Fig. 4B) suggested that non-WDV lineages indeed incorporated immune escape mutations from preceding WDV lineages. This validates our hypothesis and explains the convergence in KEScape and MLAEP scores between WDV and non-WDV lineages.

**Fig. 4:**
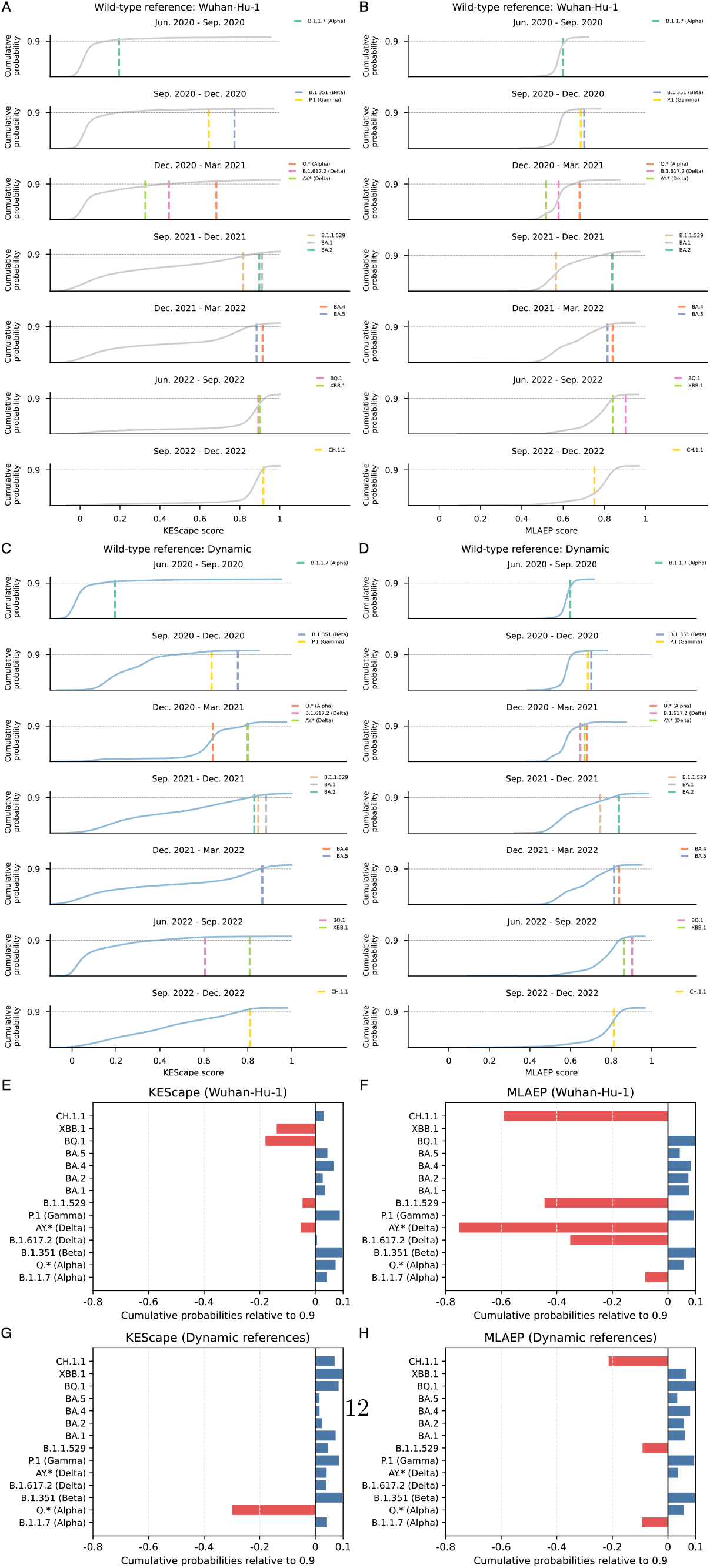
Performance of KEScape and MLAEP in prioritizing WDV lineages of SARS-CoV-2 in the lineage surveillance application. (**A-B**) Lineage surveillance evaluation using the ancestral Wuhan-Hu-1 as the wild-type reference for (**A**) KEScape and (**B**) MLAEP. The scores of non-WDV lineages are presented by cumulative distribution functions. The scores of WDV-lineages from each interval are indicated by vertical lines. (**C-D**) Lineage surveillance evaluation using dynamic references for (**C**) KEScape and (**D**) MLAEP. Data presentations of **C** and **D** follow identical conventions to **A** and **B**, respectively. (**E–H**) Predicted cumulative probabilities for WDV lineages relative to a 0.9 baseline. Predictions are shown for KEScape (**E, G**) and MLAEP (**F, H**). Wild-type references are based on the Wuhan-Hu-1 (**E–F**) or dynamic references (**G–H**).

The evolutionary trajectory of SARS-CoV-2 is predominantly characterized by continuous antigenic drift, enabling the emergence of novel lineages capable of evading host immunity established against previously prevalent lineages. To capture these dynamics, we performed an additional experiment assessing the performance using the earliest WDV lineage from the preceding interval as the wild-type reference. As shown in Fig. 4C and G, KEScape assigned scores to all but one WDV lineage that exceeded those of 90% of contemporaneous non-WDV lineages. This strong discriminatory performance is notable given the multitude of factors that influence lineage prevalence, including transmission bottlenecks [69, 70]. In contrast, MLAEP assigned three WDV lineages scores below the 90% threshold, indicating inferior performance.

We note that both KEScape and MLAEP leverage ESM2, which was pre-trained on the September 2021 release of UniRef50, a database where highly homologous viral sequences are collapsed into single representatives. In addition, ESMFold was trained on PDB chains available before May 2020, and our fine-tuned ESMFold was further trained on dozens of viral protein structures containing only two instances of the SARS-CoV-2 spike protein. Consequently, the risk of data leakage in our evaluation is effectively mitigated. Taken together, these results demonstrate that KEScape exhibits robust predictive capacity for the early identification of emerging viral lineages with immune escape potential.

### KEScape interpretability analysis and ablation study

To elucidate the internal mechanisms underlying KEScape’s capacity to model viral immune escape and evolution, we analyzed two components in the model: (i) latent embeddings preceding the final linear layer of a multilayer perceptron in the immune escape prediction step; (ii) attention weights from self-attention modules in the residue-residue dependencies identification step (Methods). Samples in the test sets of DMS datasets were used for this analysis. As demonstrated in Fig. 5A-B, SARS-CoV-2 XBB.1.5 spike and H1N1 hemagglutinin variants with immune escape capacity formed clusters that were distinct from the majority of their non-escape counterparts in the two-dimensional t-SNE projection [71], providing a robust basis for linear classification. A similar separation pattern was observed in the latent embeddings of other viral proteins (Supplementary Figs. 5-13), indicating the ability of KEScape to discriminate between immune escape and non-escape variants in the embedding space. We further quantified the local enrichment of escape variants using KNN-based enrichment scores, denoted as pos-enrichment_*k*_ (Methods). These values are shown in the lower-right corners of Fig. 5A and B. In both cases, pos-enrichment_*k*_ exceeded 1, suggesting that the model tends to embed escape variants in proximity to other escape variants.

**Fig. 5:**
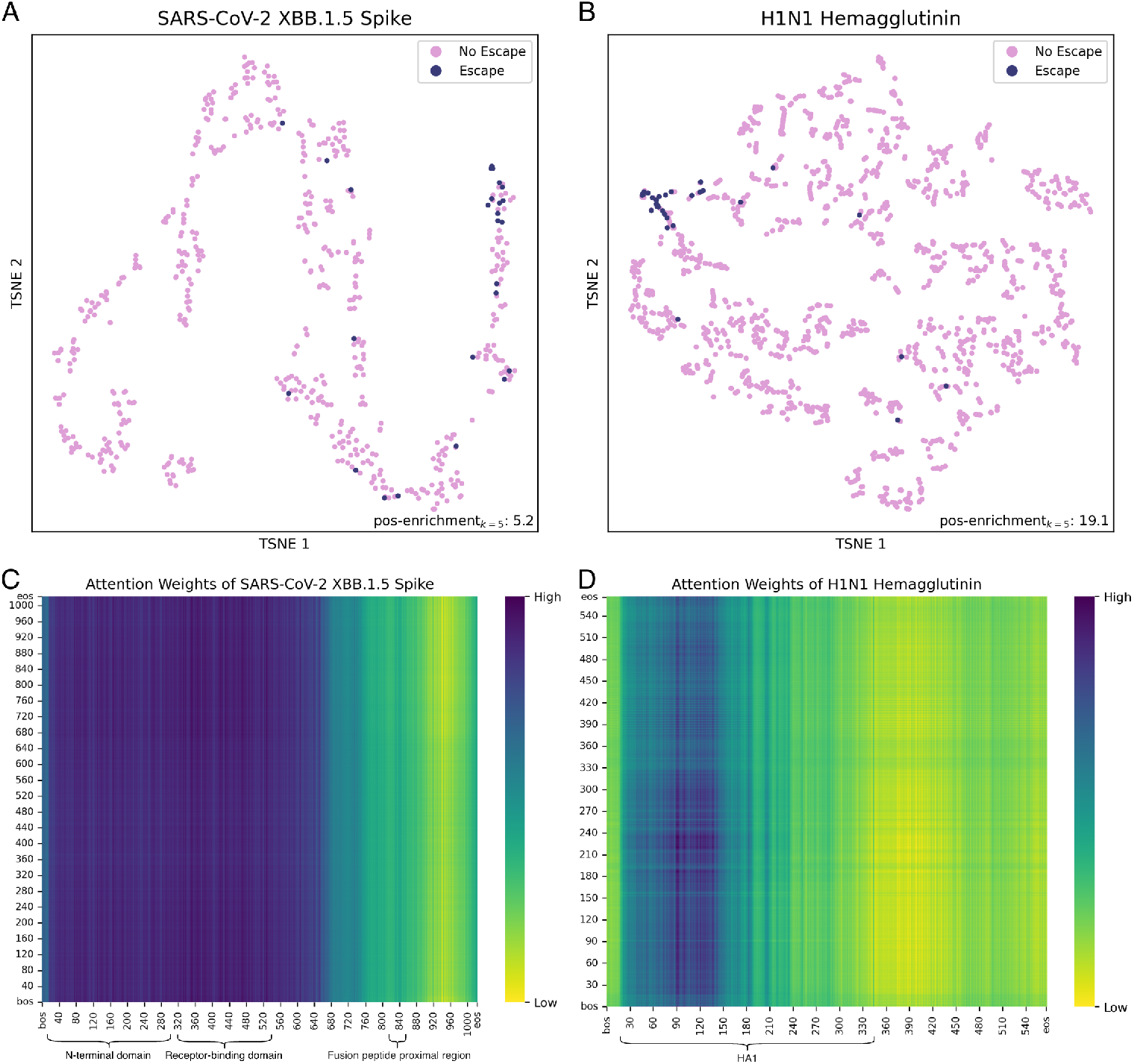
Visualization of latent embeddings and attention weights in KEScape. (**A-B**) t-SNE visualization of latent embeddings preceding the final linear layer of a multilayer perceptron in the immune escape prediction step for (**A**) SARS-CoV-2 XBB.1.5 spike and (**B**) H1N1 hemagglutinin. The pos-enrichment_*k*=5_ value is shown in the lower-right corner of each subplot. (**C-D**) Heatmaps of self-attention weights for (**C**) SARS-CoV-2 XBB.1.5 spike and (**D**) H1N1 hemagglutinin. Color intensity is proportional to attention weight magnitude, with darker hues indicating higher values.

Analysis of attention weights for SARS-CoV-2 spike (Fig. 5C) revealed pronounced attention focusing on specific tokens, with mild variation across input queries. The token with high attention weights localized to antibody-targeted regions including RBD, NTD and fusion peptide proximal region [19, 72]. A parallel analysis of H1N1 hemagglutinin similarly demonstrated focused attention on tokens within major neutralizing antibody-binding regions, particularly the HA1 subunit (Fig. 5D). The HA1 subunit, containing the receptor-binding sites and comprising the globular head, represents the primary target for neutralizing antibodies, while the HA2 subunit is occasionally targeted by neutralizing antibodies [15]. Collectively, these findings indicate that KEScape prioritizes functionally critical regions governing immune escape.

To evaluate the contribution of supervised *L*_2_ margin loss and structural embeddings, we implemented two ablated variants in which these components were removed individually. Because the supervised *L*_2_ margin loss is designed to facilitate the ranking of high-impact mutations in the context of multiple mutations, it is not expected to have a substantial effect in the DMS benchmarks, which consist only of single-mutation samples. As demonstrated in Fig. 6A-C, incorporating the supervised *L*_2_ margin loss indeed did not compromise performance on the DMS benchmarks and instead provided a marginal improvement. For the ablated variant excluding structural embeddings, KEScape outperformed this variant across the eleven DMS datasets (Fig. 6A), as evidenced by a significant increase in AUPRC. Comparative analysis of the latent embeddings for H1N1 hemagglutinin (Fig. 6B) showed that KEScape produced a more compact distribution of immune escape variants than the ablated variant without structural embeddings. This observation was further supported by the pos-enrichment_*k*=5_ values shown in the figure. We further summarized the average pos-enrichment_*k*_ across the eleven DMS datasets for *k* = 2, 5, 10 (Fig. 6C). KEScape achieved the highest average pos-enrichment_*k*_ for all three values of *k*. In addition, we investigated the contribution of these two components in the emerging lineage surveillance application. As shown in Fig. 6D, the ablated variant without supervised *L*_2_ margin loss assigned five out of fourteen WDV lineages scores below those of 90% of non-WDV lineages, whereas the variant without structural embeddings assigned three WDV lineages below this threshold. By comparison, KEScape assigned only one WDV lineage below the 90% threshold. Collectively, these results suggest that incorporating supervised *L*_2_ margin loss and structural embeddings improves the ability of KEScape to surveil emerging lineages and identify viral variants with immune escape potential. We further assessed the performance enhancement conferred by top-K *L*_2_-differential pooling. As the pooling was designed for multiple mutations while the eleven DMS datasets all consisted of single mutations, we evaluated the pooling in the emerging lineage surveillance application, utilizing the earliest WDV lineage from the preceding interval as the wild-type reference. As illustrated in Fig. 6E, top-K *L*_2_ differential pooling enabled accurate identification of predominant WDV lineages through elevated scoring relative to contemporaneous non-WDV lineages, significantly outperforming both max and mean pooling approaches. This can be attributed to that max pooling’s emphasis on the mutation with maximal embedding distance predisposes it to neglect combinatorial mutation effects and makes it more sensitive to outliers.

**Fig. 6:**
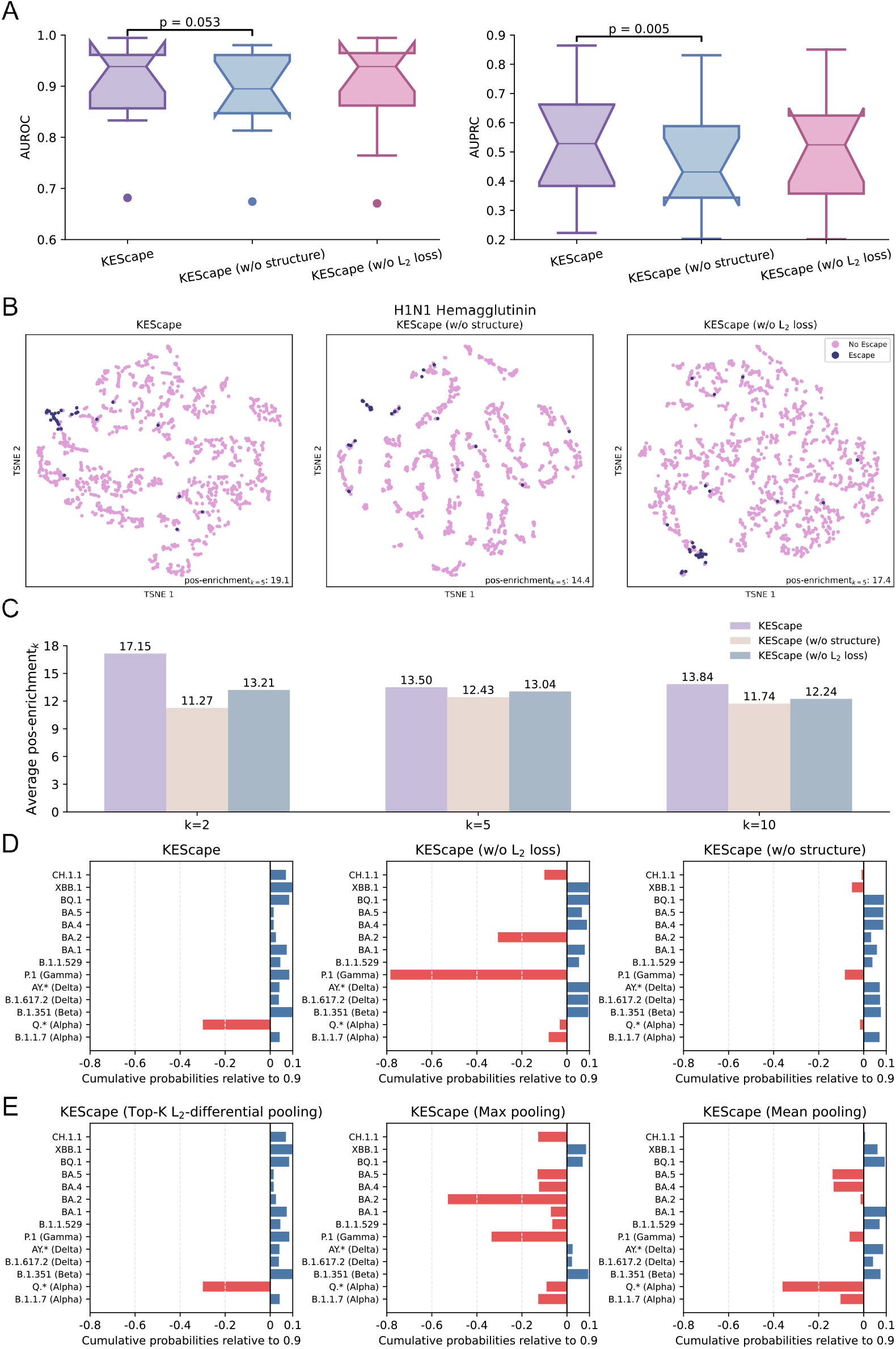
Ablation analysis of KEScape components. (**A**) Performance comparison 16 between KEScape and its ablated variants (lacking structural embeddings or the supervised *L*_2_ margin loss) across the eleven DMS datasets. The *p*-values were obtained using paired *t*-tests. (**B**) t-SNE visualization of latent embeddings preceding the final linear layer of a multilayer perceptron in the immune escape prediction step for KEScape and its ablated counterparts. (**C**) Average pos-enrichment_*k*_ (*k* = 2, 5, 10) of KEScape and its ablated counterparts. (**D-E**) Predicted cumulative probabilities relative to a 0.9 baseline for (**D**) KEScape and its ablated counterparts and (**E**) KEScape using top-K *L*_2_-differential pooling versus max pooling and mean pooling. The wild-type references are dynamic.

Conversely, the average mechanism of mean pooling can be significantly influenced by neutral mutations and thus dampen the effects of beneficial mutations due to the dominance of neutral mutations, which systematically makes mean pooling assign reduced scores for all viral variants.

## Discussion

The evolutionary arms race between viral pathogens and host immunity necessitates a paradigm shift from reactive public health measures to proactive surveillance. The efficacy of vaccines and therapeutics is perpetually challenged by antigenic drift, which allows novel variants to escape pre-existing immunity and cause successive waves of infection. To shorten the timeline for countermeasures, computational models that can accurately predict the immune escape potential of emerging variants are indispensable. These in silico approaches circumvent logistical and temporal bottlenecks of wet-lab experiments, enabling high-throughput risk assessment of viral sequences once they are detected by global surveillance systems.

Here, we present KEScape, a deep learning model that advances the state-of-the-art performance in immune escape prediction. We attribute its success to a novel architecture that synergistically integrates evolutionary context with protein structure and captures residue-residue dependencies in a sequence. Furthermore, KEScape incorporates two novel components to deal with multiple mutations: a top-K *L*_2_-differential pooling mechanism designed to account for effects from critical mutations and a supervised *L*_2_ margin loss to facilitate *L*_2_-distance-based mutation ranking. KEScape demonstrated high effectiveness by setting a new performance benchmark on DMS datasets that significantly outperformed competing models. Beyond the benchmark performance, KEScape showed a strong capacity to identify immune escape hotspots and variants and to surveil SARS-CoV-2 WDV lineages. Furthermore, KEScape exhibited a high degree of interpretability. It focused on functionally critical regions and accurately distinguished between escape and non-escape viral variants within the embedding space.

Despite these advances, wet-lab DMS experiments remain indispensable, particularly for novel viral species. As investigated in the Results, KEScape can achieve strong performance on emerging lineages that are not represented in the training set. However, this capability depends on the presence of at least a subset of training proteins from the same protein family as the target viral protein. In the absence of such condition, KEScape predictions may be unreliable. Therefore, when a novel virus emerges with pandemic potential, wet-lab DMS experiments against the wild-type virus are essential during the initial wave of large-scale infection or vaccination. Once population-level adaptive immunity has been established and the virus is subjected to strong immune-mediated selective pressure, computational models such as KEScape can be applied to surveil newly emerging variants using the DMS dataset.

An ideal predictive framework would identify high-risk variants at their genesis and accurately forecast their subsequent evolutionary trajectories. However, the inherent stochasticity of viral evolution, influenced by factors such as transmission bottlenecks, renders such deterministic prediction currently impractical. A more practical application, for which KEScape is well-suited, is to generate a prioritized pool of candidate variants with high predicted immune escape potential based on real-time genomic surveillance. This allows for the immediate computational assessment of newly detected variants, identifying high-risk candidates for rapid experimental validation and informing public health responses. While the prediction of viral immune escape remains a formidable challenge due to biological complexity and stochastic effects, the progress demonstrated here and elsewhere in the field represents a significant step towards developing the tools necessary to combat future pandemics.

## Methods

### Deep mutational scanning datasets

We collected eleven DMS datasets of various viral proteins (Supplementary Table 1). Since the three DMS datasets obtained from EVEscape were already processed, we performed data processing on the remaining eight DMS datasets. Specifically, we removed any mutations that (i) had no recorded immune escape scores, (ii) included ambiguous amino acids (i.e. X, B, Z, J), or (iii) resulted in a premature stop codon. If negative immune escape scores were present in a dataset, we applied an exponential transformation to the scores. In particular, we used base-2 exponential transformations for the SARS-CoV-2 XBB.1.5 spike, H5N1 hemagglutinin, Nipah receptor-binding protein, lassa glycoprotein, H3N2 hemagglutinin, rabies glycoprotein datasets, whereas a base-10 exponential transformation was applied to the H3N2 neuraminidase dataset. For datasets containing multiple immune escape scores per mutation from different neutralizing antibodies or sera (including lassa glycoprotein, H3N2 hemagglutinin and rabies glycoprotein), we used the maximum immune escape scores for each mutation. To mitigate noises in immune escape scores, we adopted the binarization strategy from EVEscape. For each dataset, we fitted a gamma distribution using the method of moments and set a threshold corresponding to a 5% false discovery rate. Mutations with immune escape scores greater than or equal to this threshold were labeled as conferring immune escape (label 1), whereas those below the threshold were labeled as non-escape (label 0). We excluded any mutations in the test set that could not be predicted by EVEscape due to insufficient coverage in its multiple sequence alignment. This ensured consistent test sets across all computational models.

For the SARS-CoV-2 DMS dataset obtained from E2VD, we constructed a transformed dataset that used a single immune escape score for each mutation by selecting the maximum value observed across all tested neutralizing antibodies. This transformed dataset was then randomly divided into training (70%), validation (15%) and test (15%) sets, with the constraint that no mutation appeared in more than one subset. The original, non-transformed dataset was subsequently partitioned into training, validation, and test sets by mapping each sample according to the split assignment of its corresponding mutation in the transformed dataset. The test set was filtered to be consistent with the transformed one by selecting only the antibody with the largest immune escape score for each mutation. For binarization, we set the threshold to 0.4, consistent with E2VD. We trained E2VD using its default settings on the training set and evaluated its performance on the test set. KEScape was trained on the transformed training set, and all models except E2VD were evaluated on the transformed test set.

### EVEscape and CSCS fine-tuning

We included fine-tuned versions of EVEscape and CSCS in the benchmark. For EVEscape, we used four features as input to a multilayer perceptron: the embedding from the final hidden layer of the EVE encoder before the mean and log-variance heads for the mutant sequence (*H*_*e*_), the EVE score (*S*_*EVE*_), the weighted contact number (*S*_*WCN*_), and the dissimilarity score (*S*_*D*_). Specifically, the fine-tuned EVEscape computes the immune escape score of a mutant sequence as follows:

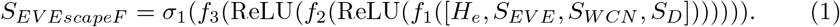

Here, *f*_1_, *f*_2_, and *f*_3_ denote linear transformations; ReLU denotes the rectified linear unit activation function; [*H*_*e*_, *S*_*EVE*_, *S*_*WCN*_, *S*_*D*_] denotes the concatenation of the four features; and *σ*_1_ denotes the sigmoid function. The model was fine-tuned using a weighted cross-entropy loss function with a learning rate of 0.001.

For CSCS, we initialized the model from the protein-specific pre-trained checkpoint and appended an immune escape prediction head. This head takes as input the final hidden state of the LSTM encoding the left sequence context (*H*_*l*_) and the final hidden state of the LSTM encoding the right sequence context (*H*_*r*_). These two context embeddings are concatenated and passed through a multilayer perceptron, analogous to the fine-tuning head used for EVEscape, to produce immune escape scores:

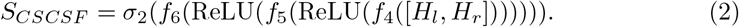

In this expression, *f*_4_, *f*_5_, and *f*_6_ denote linear transformations; ReLU denotes the rectified linear unit activation function; [*H*_*l*_, *H*_*r*_] denotes the concatenation of *H*_*l*_ and *H*_*r*_; and *σ*_2_ denotes the element-wise sigmoid function. The prediction head outputs a vector of scores over amino acid identities conditioned on the sequence context surrounding the target residue. During training, the score corresponding to the mutant amino acid at the target position is selected and optimized against the ground-truth label using cross-entropy loss with a learning rate of 0.001. As a result, only substitutions observed in the training set contribute directly to the loss, whereas substitutions absent from the training set do not receive direct supervision.

### Immune escape hotspot experiment

Given twenty different amino acids, with one wild-type and nineteen alternative amino acids for a residue *i*, the max per-residue KEScape score *m*^(*i*)^ is defined to be

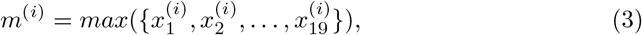

where 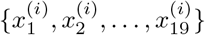 is a set of all immune escape scores of the mutant sequences with alternative amino acids. Similarly, the mean per-residue KEScape score *z*^(*i*)^ is defined to be

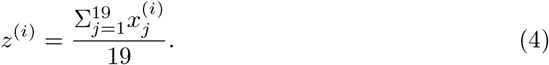

The max per-residue KEScape scores were calculated and mapped onto a viral variant of the SARS-CoV-2 BA.2 lineage (pdb: 7XIX).

### SARS-CoV-2 lineage surveillance dataset

We used a dataset from EVEscape, comprising viral sequences from GISAID sampled across diverse geographical regions over time. Based on the WHO and PANGO nomen-clature [73], we classified lineages associated with WHO-designated variants (referred to as WDV lineages), as listed in Supplementary Table 2. All remaining lineages were classified as non-WDV lineages. The dataset aggregated viral isolates with the same spike protein sequences into a group. For each group, the earliest collection date among all constituent isolates was assigned as the representative date for that group. In addition, a definitive PANGO lineage was assigned to each group: if any isolate within a group belonged to WDV lineages, the entire group was classified under that lineage. Groups were subsequently filtered to exclude any that contained ambiguous amino acids, premature stop codons, or were represented by ten or fewer viral isolates (count ≤ 10). To reflect the dynamics of viral lineage turnover, we divided the timeline into non-overlapping 90-day intervals. WDV lineages present in previous intervals were excluded from subsequent intervals, and any intervals without newly emerged WDV lineages were also removed. We retrieved immune escape mutations from Alam et al. [68], calculated the proportion of these mutations in non-WDV lineages over different intervals, and investigated their presence in WDV lineages (Supplementary Figure 4). We conducted two experiments with different wild-type references: one using the ancestral Wuhan-Hu-1 (a constant reference) and one using the earliest WDV lineage from the preceding interval (a dynamic reference).

The immune escape potential of each unique viral spike sequence was predicted by KEScape relative to both the constant and dynamic references. Since a WDV lineage can encompass multiple variants with different sets of mutations within a time interval, a representative score was required. Therefore, to quantify the immune escape potential of a WDV lineage, the maximum KEScape score among all its constituent variants within that interval was selected.

### KEScape architecture

KEScape is designed to quantify the immune escape potential of a given viral variant. The model takes a mutant sequence, *S*_*m*_, and its corresponding wild-type reference sequence, *S*_*w*_, as input. It computes a score, denoted as 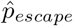, which is constrained to the interval (0, 1). A higher score signifies an increased probability that the viral variant will evade host immune surveillance. This predictive process can be mathe-matically formulated by representing the KEScape model as a function *f*_*KEScape*_, such that:

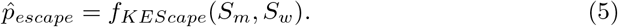

The inference process of KEScape comprises four primary steps. KEScape begins with the generation of protein embeddings from a fine-tuned ESMFold (650M parameter version, see supplementary notes), denoted as *f*_*ESMFold*_. For a given wild-type and mutant protein sequence pair, the model processes each sequence respectively. Input sequences exceeding a length of 1022 amino acids should be truncated to meet the model’s architectural constraints. ESMFold generates two distinct sets of embeddings for each residue in a sequence: evolutionary embeddings and structural embeddings. Specifically, for a wild-type sequence *S*_*w*_ of length *L*_*w*_, this operation yields evolutionary embeddings 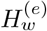 with dimensions (*L*_*w*_ +2, 1280), and structural embeddings 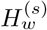 with dimensions (*L*_*w*_, 1024); similarly, An analogous procedure is applied to the mutant sequence *S*_*m*_ to produce its respective embeddings, 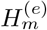 and 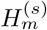. This process can be formally expressed as:

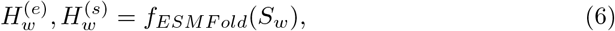

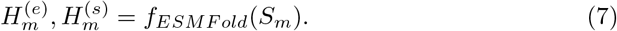

Here, 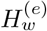 represents the set of evolutionary embeddings 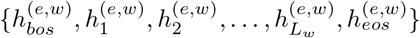, including embeddings for each residue, augmented by *bos* and *eos* tokens. In contrast, 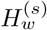 is the set of structural embeddings 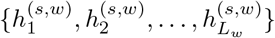, corresponding only to the residues of the sequence. The embeddings for the mutant sequence, 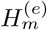 and 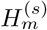 are generated analogously.

In the second step, KEScape integrates evolutionary embeddings and structural embeddings into unified embeddings for each sequence. For clarity, the following procedure is described for the wild-type sequence. The dimensionality of the evolutionary embeddings 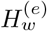 is first reduced from 1280 to 1024 via a linear projection layer *f*_*linear*1_ to match the one from the structural embeddings.

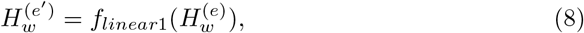

where the resulting 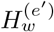 has dimensions of (*L*_*w*_ + 2, 1024). These projected evolutionary embeddings are transformed to queries, and the structural embeddings, 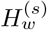, are transformed to keys and values as follows:

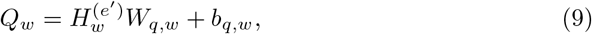

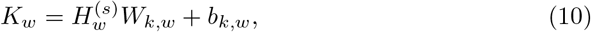

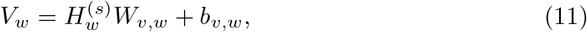

where *W*_*q,w*_, *W*_*k,w*_, *W*_*v,w*_ are the weights and *b*_*q,w*_, *b*_*k,w*_, *b*_*v,w*_ are the biases. *Q*_*w*_, *K*_*w*_, and *V*_*w*_ are subsequently divided into 8 heads as 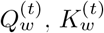 and 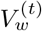 for a given head *t*. Subsequently, 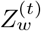, the weighted sum of the values is calculated as follows:

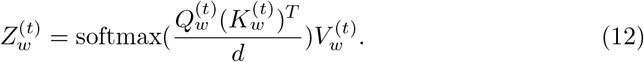

In this formulation, *d* is a scaling factor, and 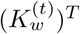 is the transpose of 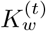. The output matrices from all eight heads are concatenated and passed through a linear projection (*f*_*linear*2_) to produce a representation *Z*_*w*_:

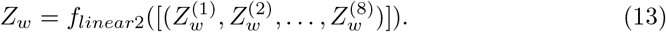

The resulting *Z*_*w*_ has the dimensions of (*L*_*w*_ + 2, 1024). Finally, this representation is processed through a multilayer perceptron with residual connections to yield unified embeddings for the wild-type sequence. The unified embeddings for the mutant sequence can be derived analogously.

Following the integration of structural embeddings and evolutionary embeddings, KEScape employs a self-attention module to further refine the unified embeddings by modeling residue-residue dependencies. For each sequence, the unified embeddings are projected into query *Q*, key *K* and value *V*. The self-attention module then computes a contextually updated representation, *Z*^′^, by calculating a weighted sum of the values, where the weights are determined by the similarity between queries and keys:

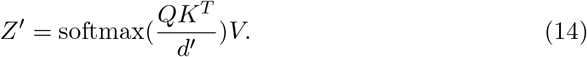

Here, *d*^′^ is a scaling factor. This output is subsequently processed by a multilayer perceptron with residual connections, yielding the context-aware unified embeddings for each sequence.

In the fourth step, KEScape employs a top-K *L*_2_-differential pooling mechanism to identify and select mutations with large *L*_2_ distances. For a mutant sequence with *N* mutations, the *L*_2_ distance is calculated between the wild-type and mutant context-aware unified embeddings at each of the *N* mutant positions. Suppose the corresponding *N* embeddings for the wild-type and mutant sequence are 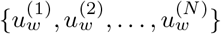 and 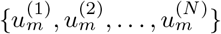, respectively, the *L*_2_ distance at a given mutant position *i* is calculated as

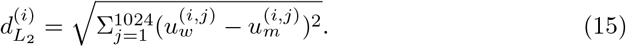

Here, 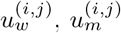 are the *j*-th elements of the vector 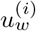 and 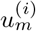, respectively. The top-*K* mutant positions are selected by ranking the *L*_2_ distances in descending order. Let

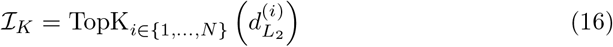

denote the index set of the *K* mutant positions with the largest *L*_2_ distances, where |ℐ |_*K*_ = *K*. The corresponding wild-type and mutant context-aware unified embeddings are then separately averaged to obtain the pooled embeddings:

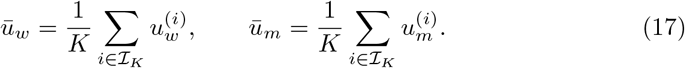

KEScape then constructs a representation by concatenating the wild-type pooled embedding, the mutant pooled embedding, and their difference:

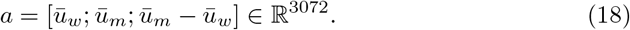

This representation is passed through a multilayer perceptron consisting of two linear layers with a GELU activation between them, followed by a softmax layer:

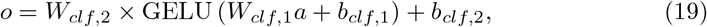

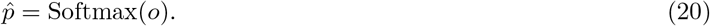

Here, *W*_*clf*,1_ and *b*_*clf*,1_ are the weight matrix and bias vector of the first linear layer, *W*_*clf*,2_ and *b*_*clf*,2_ are those of the second linear layer, and *o* denotes the output logits. The final vector 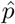 represents the predicted class probabilities, with the immune escape probability given by

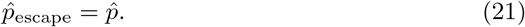

### Supervised *L*_2_ margin loss

Supervised *L*_2_ margin loss is designed to explicitly force the model to increase the *L*_2_ distances between the wild-type and mutant embeddings for immune escape variants while decreasing such distances for non-escape variants in the embedding space. For each sample, the average *L*_2_ distances across the selected *K* mutations is defined as:

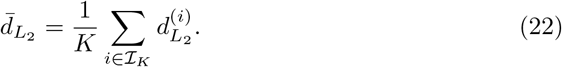

During training, each sample is associated with a ground-truth label *y* ∈ {0, 1}, where *y* = 1 denotes an immune escape variant (positive class) and *y* = 0 denotes a non-escape variant (negative class). The supervised *L*_2_ margin loss applies distinct penalties based on the class label. For positive samples, the model is penalized if 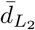 is smaller than a predefined positive margin *m*_*pos*_. Conversely, for negative samples, the model is penalized if 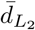 exceeds a predefined negative margin *m*_*neg*_. To prevent extreme penalties from dominating the gradient, the penalties are capped by a ceiling parameter *C*.

The individual sample penalties 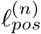 and 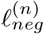 for a given sample *n* are formulated as

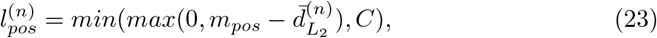

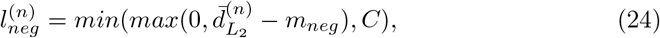

for positive and negative classes, respectively, where 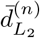 denotes the mean *L*_2_ distance for sample *n*.

For a given training batch, let 𝒫 denote the set of indices for positive samples (*y* = 1) and 𝒩 denote the set of indices for negative samples (*y* = 0), the supervised *L*_2_ margin loss is then defined as:

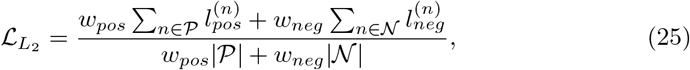

where *w*_*pos*_ and *w*_*neg*_ are the class weights for positive and negative samples, respectively, and |𝒫| and |𝒩| are the number of samples in each class within the batch.

### KEScape training

KEScape was trained using a combined loss function consisting of weighted binary cross-entropy loss and supervised *L*_2_ margin loss. The optimizer was selected as AdamW [74] with *β*_1_ = 0.9, *β*_2_ = 0.98, and a weight decay of 0.01. The gradient accumulation was employed to achieve an effective batch size of 128. A learning rate scheduler with a warm-up phase was implemented. The learning rate increased to a peak value over the first *min*(*int*(0.05 ∗ *num training steps*), 500) steps, and then decayed to a final value of 10% of its peak value in the subsequent training period. Each DMS dataset was partitioned into training (70%), validation (15%), and test (15%) sets. In the DMS benchmark experiments, KEScape was trained separately on the training set of each dataset and evaluated on the corresponding test set using the checkpoint that achieved the lowest cross-entropy loss on the validation set. The peak learning rate was set to 0.001. In zero-shot experiments, including the identification of immune escape hotspots and variants from emerging lineages as well as lineage surveillance, KEScape was trained on those DMS datasets that excluded the target lineages, with a peak learning rate of 0.0005. The only exception was the identification of immune escape hotspots in the BA.2 spike protein, for which KEScape was trained on all DMS datasets except the XBB.1.5 spike dataset (There is no BA.2 spike DMS dataset in the benchmark). For the comparative benchmark against E2VD, training was restricted to the SARS-CoV-2 DMS dataset provided by E2VD, with training, validation, test splits of 70%, 15%, 15%, respectively.

### Model interpretation and ablation study

To elucidate the decision-making process of KEScape, we conducted model interpretation experiments. First, we analyzed the latent embeddings preceding the final linear layer of a multilayer perceptron in the immune escape prediction step. These highdimensional embeddings were projected into a two-dimensional space using t-SNE to visualize the distribution of immune escape and non-escape viral variants. In addition, we used KNN-based positive enrichment score to quantify the degree of enrichment for embeddings of escape variants in the high-dimensional embedding space. The score is defined as follows:

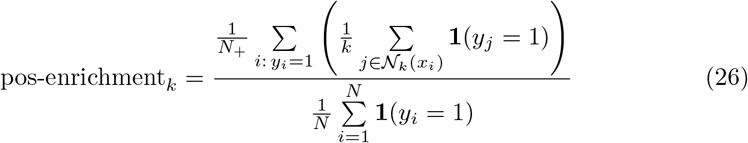

where *N* is the total number of samples, 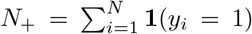 is the number of positive samples, 𝒩_*k*_(*x*_*i*_) denotes the set of *k*-nearest neighbors of sample *x*_*i*_, *y*_*i*_ ∈ {0, 1} is the class label of sample *i*, and **1**(*y*_*j*_ = 1) is the indicator function equal to 1 if neighbor *j* is positive and 0 otherwise.

Second, we examined pre-softmax attention weights from the self-attention modules, which are responsible for identifying residue-residue dependencies. These weights were visualized as heatmaps to identify the specific residue-residue interactions that the model weighted most heavily in its computations.

To evaluate the contribution of integrating the supervised *L*_2_ margin loss and structural embeddings, an ablation study was conducted wherein these two components were removed respectively. Specifically, for the ablated variant excluding the supervised *L*_2_ margin loss, this loss was removed from the loss function while the others remained the same. For the ablated variant excluding the structural embeddings, the corresponding structural embeddings input to the cross-attention module were replaced with a duplicate of the evolutionary embeddings. Accordingly, the linear layer in the evolution-structure fusion block was applied to both evolutionary embeddings to ensure dimensional compatibility. In a separate ablation study targeting the top-K *L*_2_-differential pooling mechanism, we replaced the mechanism with max pooling and mean pooling, respectively. Max pooling was defined as selecting a single mutation with the largest *L*_2_ distance between its wild-type and mutant context-aware unified embeddings. In contrast, mean pooling involved averaging the context-aware unified embeddings across all mutant positions for the wild-type and mutant sequences, respectively.

## Data availability

The sources of eleven DMS datasets are provided in Supplementary Table 1 and these datasets are available at https://github.com/ericcombiolab/KEScape. The DMS dataset from E2VD can be downloaded from https://github.com/ZhiweiNiepku/E2VD. The SARS-CoV-2 BA.2 spike protein structure (pdb: 7XIX) is available at https://www.rcsb.org/structure/7XIX. The SARS-CoV-2 lineage surveillance dataset is available at https://marks.hms.harvard.edu/evescape/strain scores 20230318.zip.

## Code availability

The source code of KEScape is available at https://github.com/ericcombiolab/KEScape.

## Acknowledgements

The project is partially supported by the Young Collaborative Research grant (No. C2004-23Y) and HMRF grant (No. 11221026). Fig. 1A was created with BioRender. com.

## Author contributions

C.W. and L.Z. conceived the study. C.W. developed the methods and conducted the experiments. L.Z. contributed to technical discussions. C.W. wrote the manuscript. L.Z. revised the manuscript.

## Supplementary information

A supplementary file is attached.

