## supplementary file for "A structure-informed evolutionary model for predicting viral immune escape and evolution"

#### Contents

**1 Supplementary notes** **1**
1.1 Top-K $L_2$-differential pooling . . . . . . . . . . . . . . . . . . . . . 1
1.2 ESMFold fine-tuning . . . . . . . . . . . . . . . . . . . . . . . . . . . 1

**2 Supplementary Tables** **2**

**3 Supplementary Figures** **3**

#### 1 Supplementary notes

##### 1.1 Top-K $L_2$-differential pooling

The algorithm of the top-K $L_2$-differential pooling mechanism is described below:

**Algorithm 1** Top-K $L_2$-Differential Pooling

**def** topKL2DifferentialPooling($\{X^{wild}\}$, $\{X^{mut}\}$, $N$, $K$) **:** $\#X^{wild}.shape = (N, embed\_dim), X^{mut}.shape = (N, embed\_dim)$

$K = min(N, K)$ # deal with the cases when N is smaller than K
$distances = L2norm(X^{mut} - X^{wild}, dim = 1)$
$ti = fetch\_K\_largest\_dist\_indices(distances, num = K)$
$x\_wild = mean(X^{wild}_{ti}, dim = 0)$
$x\_mut = mean(X^{mut}_{ti}, dim = 0)$
**return** $x\_wild, x\_mut$

##### 1.2 ESMFold fine-tuning

KEScape incorporates the 650M parameter version of the ESMFold. As initial evaluations of the original ESMFold exhibited suboptimal performance in the target viral proteins, we fine-tuned the ESMFold with dozens of viral proteins and human receptor proteins whose structures were generated by Chai-1 [1]. The fine-tuning process was guided by a composite loss function composed of frame aligned point error (FAPE) loss, distogram loss, and violation loss. During the fine-tuning, the ESM-2 parameters were frozen, with updates applied exclusively to the structure modules in ESMFold. Optimization was performed using the Adam optimizer [2] with a weight decay of 0.01. A learning rate scheduler was employed, which increased the learning rate to a peak value of $8 \times 10^{-6}$ over the first 1200 steps, and then finally decayed to one tenth of its peak value in the subsequent training period.

#### 2 Supplementary Tables

| DMS dataset | Reference | Website |
|---|---|---|
| SARS-CoV-2 spike | [3] | SARS-CoV-2 |
| SARS-CoV-2 XBB.1.5 spike | [4] | SARS-CoV-2 XBB |
| H1N1 hemagglutinin | [3] | H1N1 HA |
| HIV envelope | [3] | HIV envelope |
| H5N1 hemagglutinin | [5] | H5N1 HA |
| Zika envelope | [6] | Zika envelope |
| Nipah receptor-binding protein | [7] | Nipah RBP |
| Lassa glycoprotein | [8] | Lassa glycoprotein |
| H3N2 hemagglutinin | [9] | H3N2 HA |
| H3N2 neuraminidase | [10] | H3N2 NA |
| Rabies glycoprotein | [11] | Rabies glycoprotein |

Table 1: The reference and website for each DMS dataset.

| PANGO nomenclature | WHO nomenclature |
|---|---|
| B.1.1.7 | Alpha |
| Q.* | Alpha |
| B.1.351 | Beta |
| B.1.617.2 | Delta |
| AY.* | Delta |
| P.1 | Gamma |
| B.1.1.529 | Omicron |
| BA.1 | Omicron |
| BA.2 | Omicron |
| BA.4 | Omicron |
| BA.5 | Omicron |
| BQ.1 | Omicron |
| XBB.1 | Omicron |
| CH.1.1 | Omicron |

Table 2: The mapping of PANGO nomenclature and WHO nomenclature for WDV lineages.

### 3 Supplementary Figures

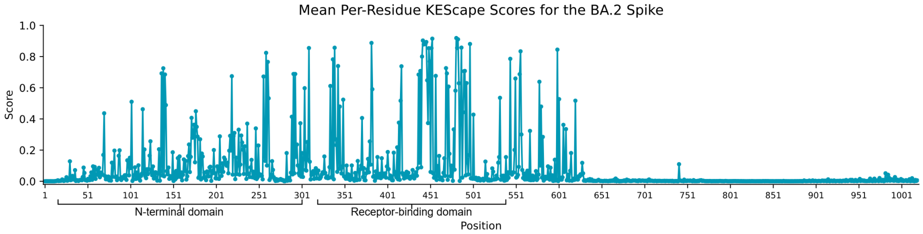


Supplementary Fig. 1: Mean per-residue KEScape scores for the BA.2 spike.

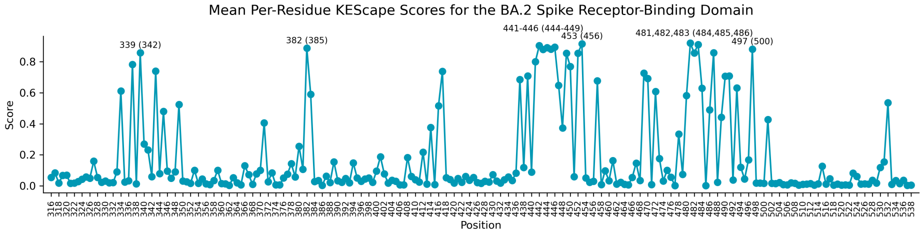


Supplementary Fig. 2: Mean per-residue KEScape scores for the RBD of the BA.2 spike. Residues with higher scores are highlighted, with the Wuhan-Hu-1 reference positions provided in parentheses.

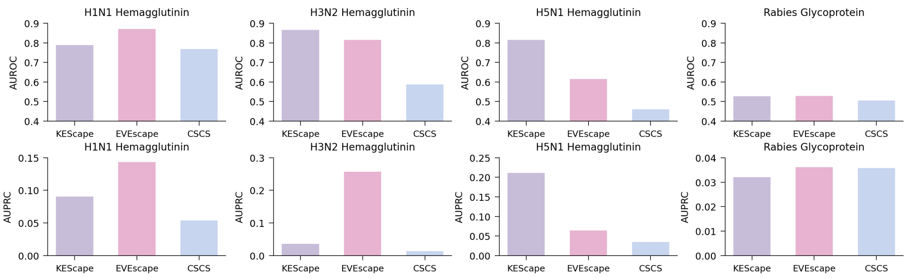


Supplementary Fig. 3: Zero-shot performance of KEScape, EVEscape, and CSCS on the H1N1 hemagglutinin, H3N2 hemagglutinin, H5N1 hemagglutinin, rabies glycoprotein DMS datasets, respectively.

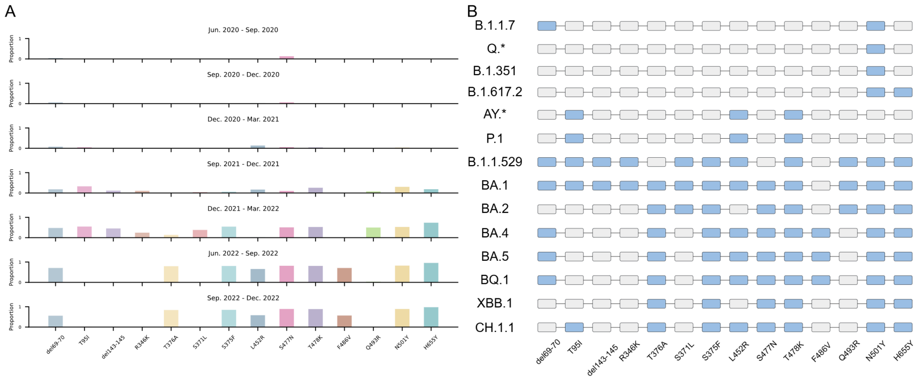


Supplementary Fig. 4: (**A**) The proportion of immune escape mutations found within non-WDV lineages over different intervals. (**B**) A matrix documenting presence (blue) versus absence (gray) of immune escape mutations across WDV lineages.

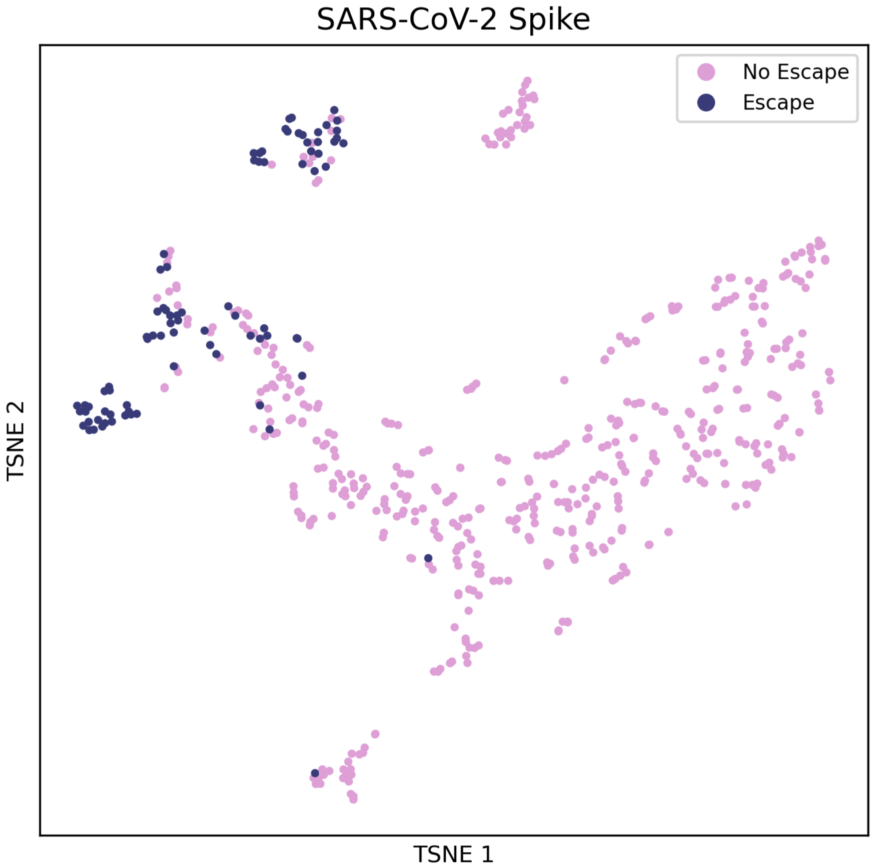


Supplementary Fig. 5: t-SNE of latent embeddings preceding the final linear layer of a multilayer perceptron in the immune escape prediction step for SARS-CoV-2 spike.

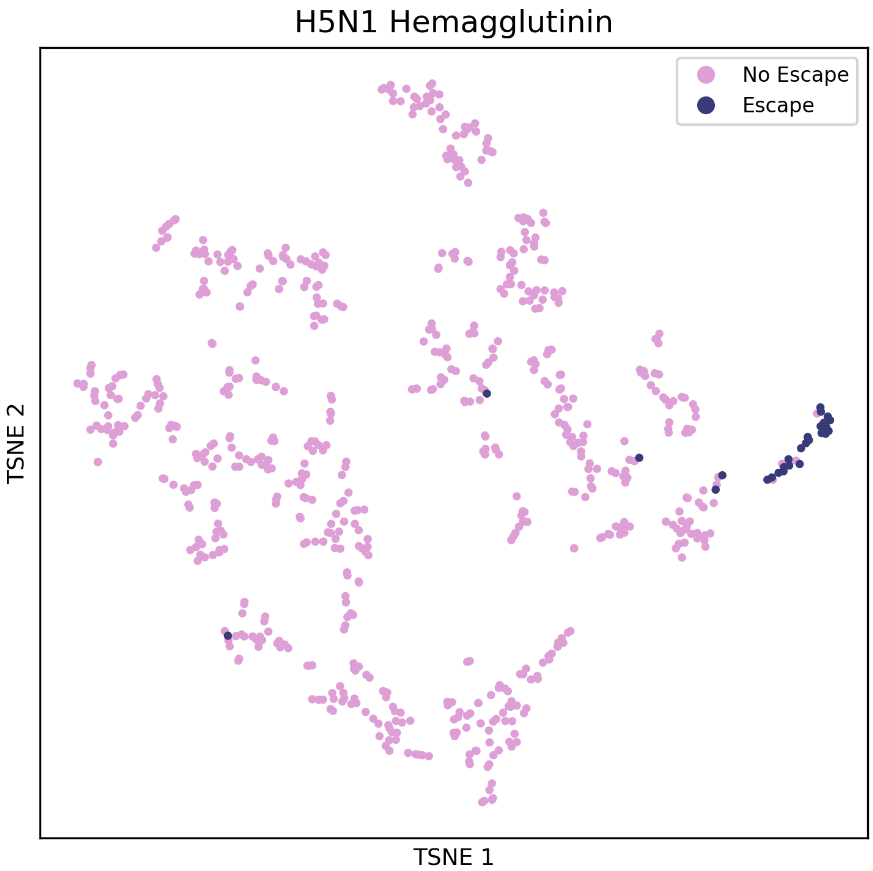


Supplementary Fig. 6: t-SNE of latent embeddings preceding the final linear layer of a multilayer perceptron in the immune escape prediction step for H5N1 hemagglutinin.

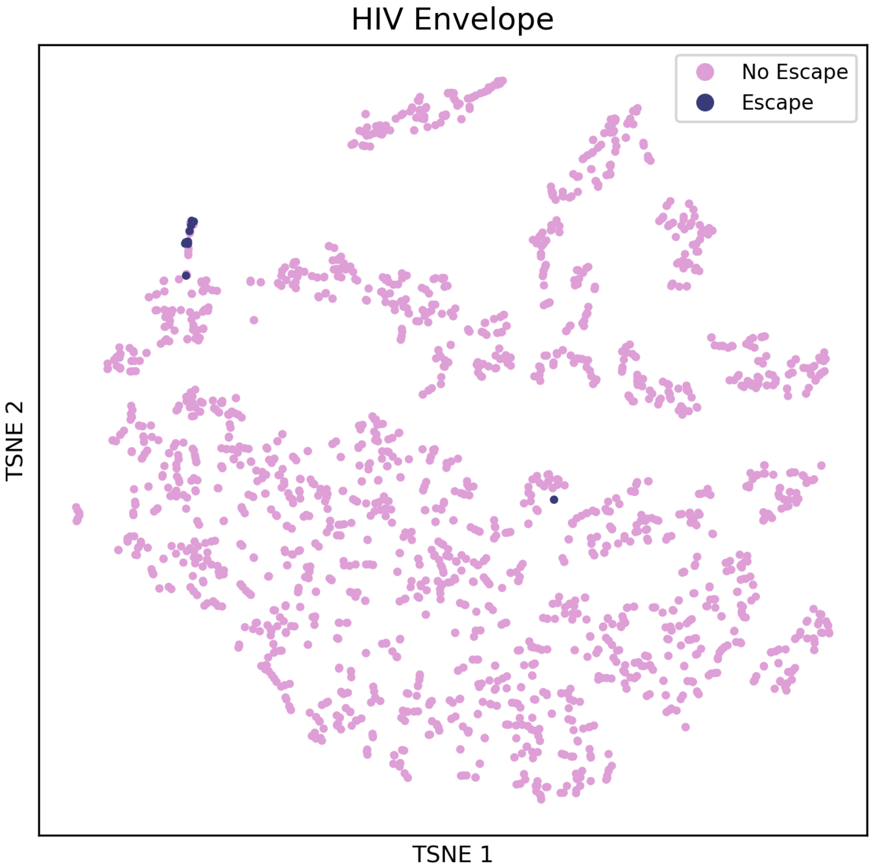


Supplementary Fig. 7: t-SNE of latent embeddings preceding the final linear layer of a multilayer perceptron in the immune escape prediction step for HIV envelope.

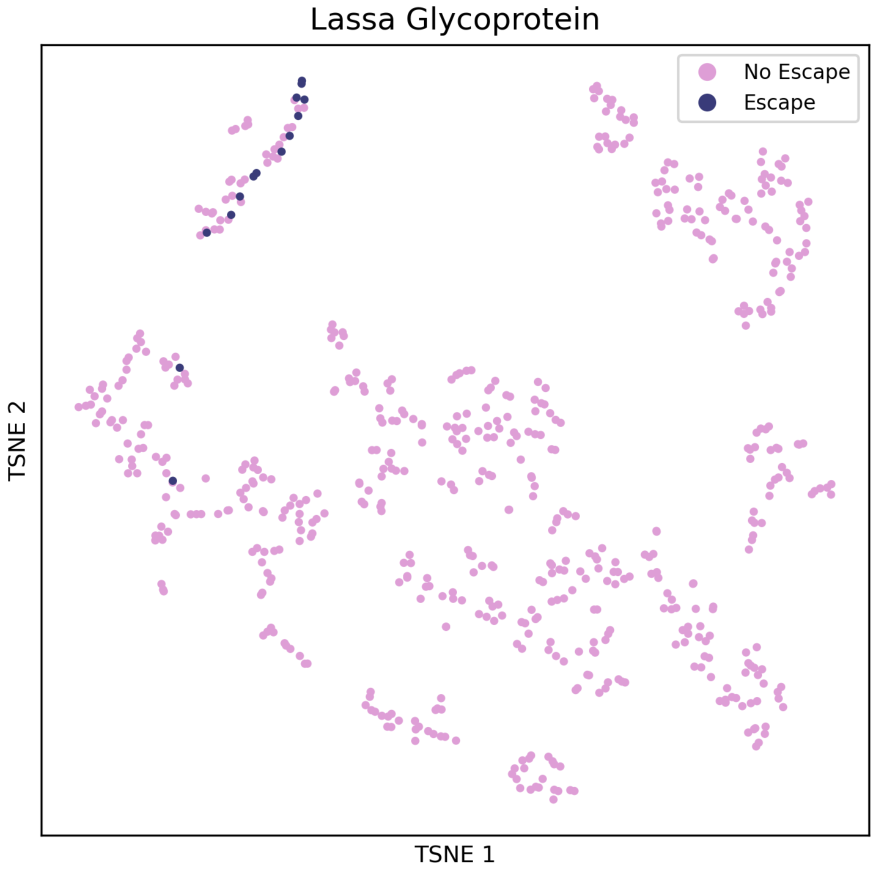


Supplementary Fig. 8: t-SNE of latent embeddings preceding the final linear layer of a multilayer perceptron in the immune escape prediction step for Lassa glycoprotein.

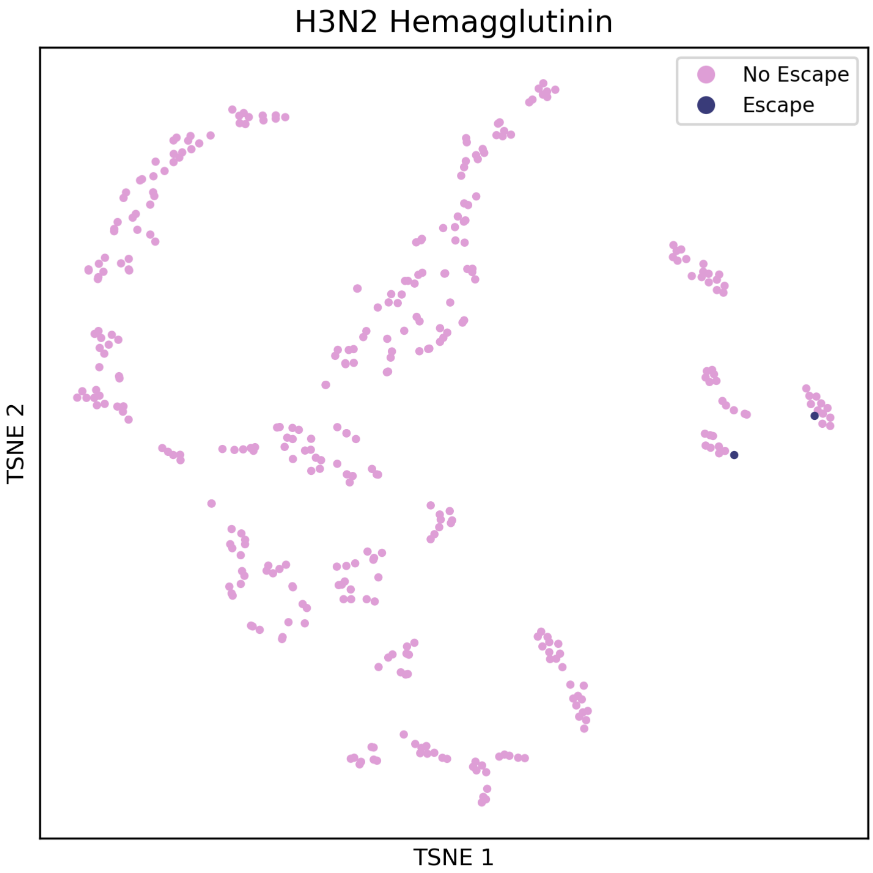


Supplementary Fig. 9: t-SNE of latent embeddings preceding the final linear layer of a multilayer perceptron in the immune escape prediction step for H3N2 hemagglutinin.

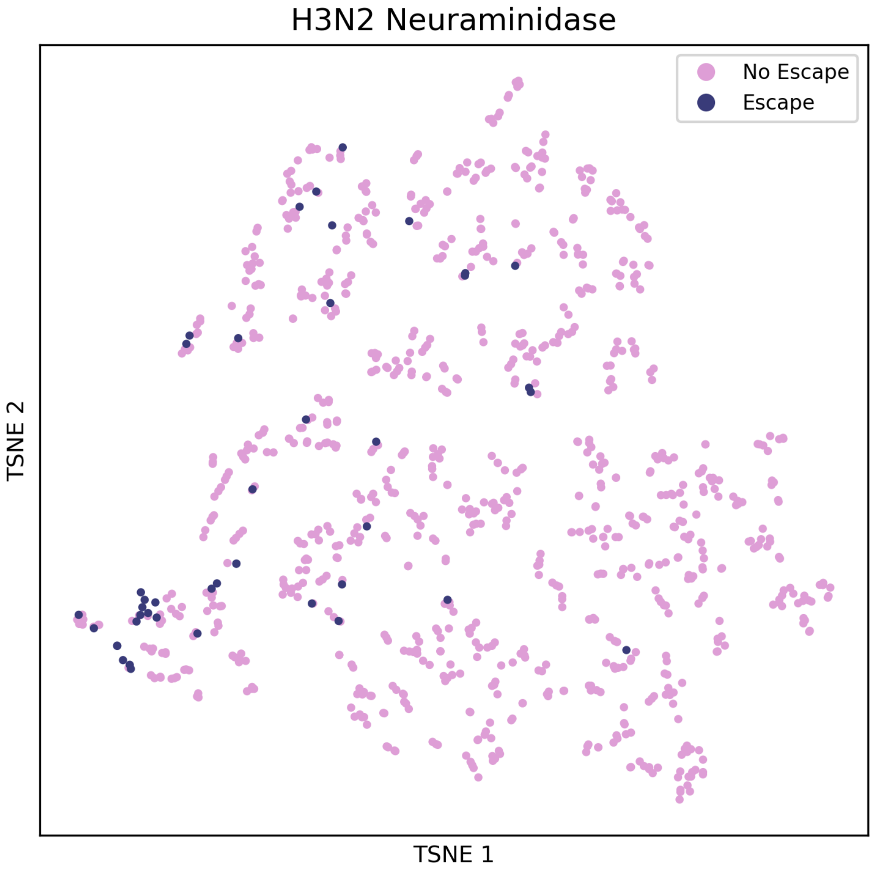


Supplementary Fig. 10: t-SNE of latent embeddings preceding the final linear layer of a multilayer perceptron in the immune escape prediction step for H3N2 neuraminidase.

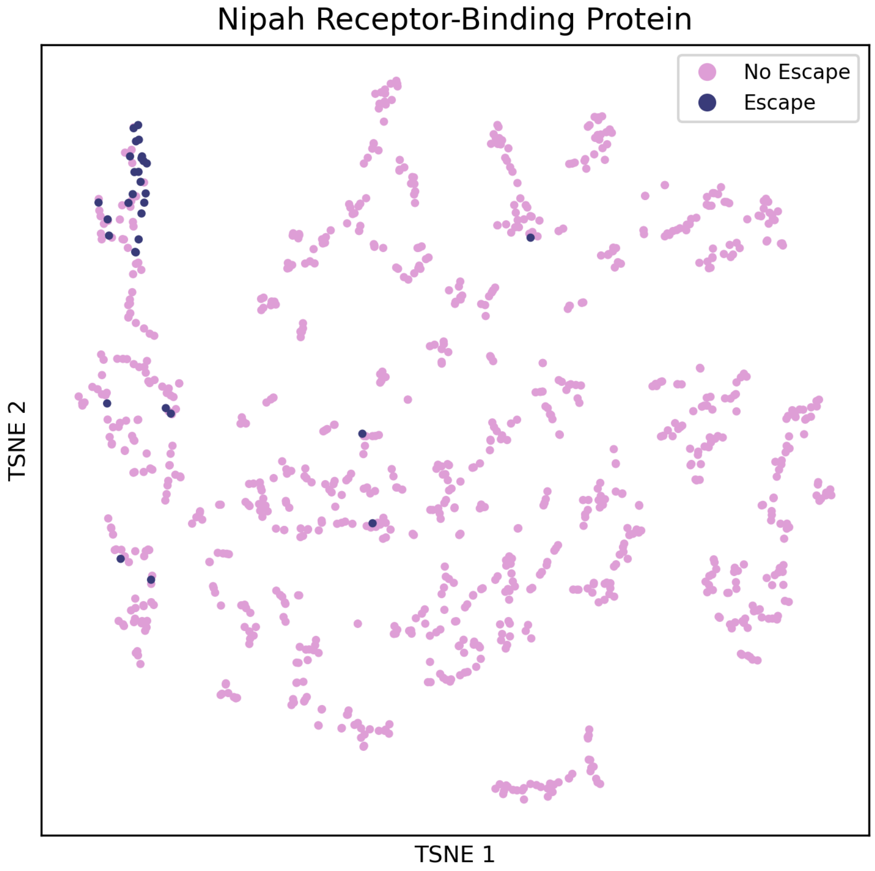


Supplementary Fig. 11: t-SNE of latent embeddings preceding the final linear layer of a multilayer perceptron in the immune escape prediction step for Nipah receptor-binding protein.

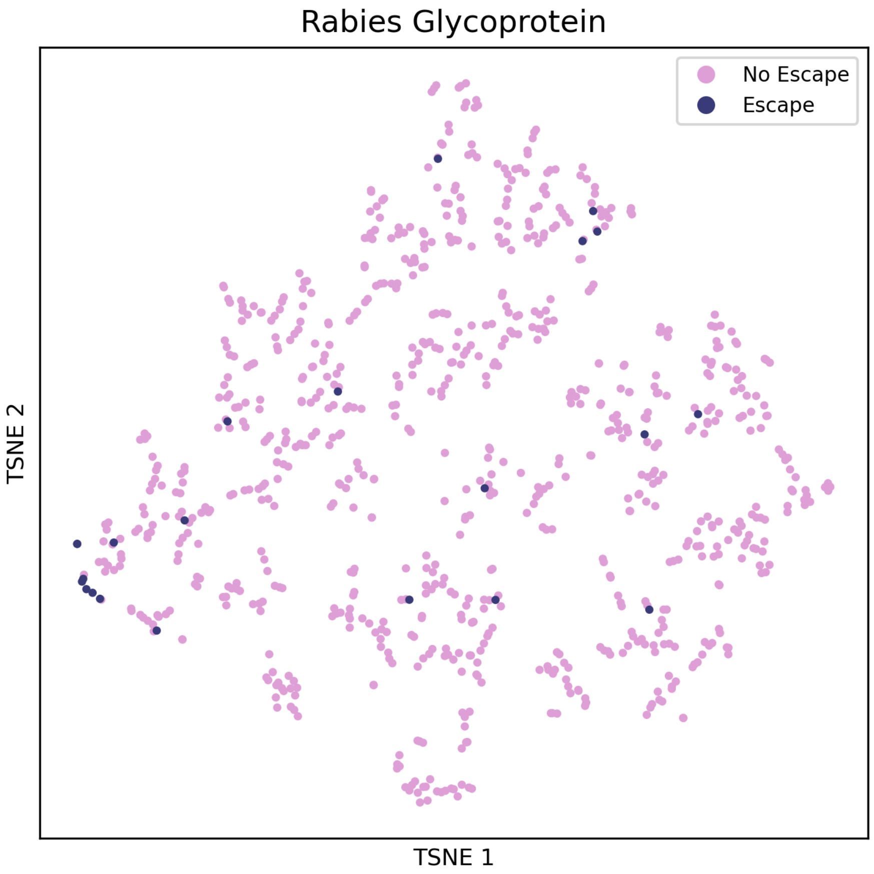


Supplementary Fig. 12: t-SNE of latent embeddings preceding the final linear layer of a multilayer perceptron in the immune escape prediction step for rabies glycoprotein.

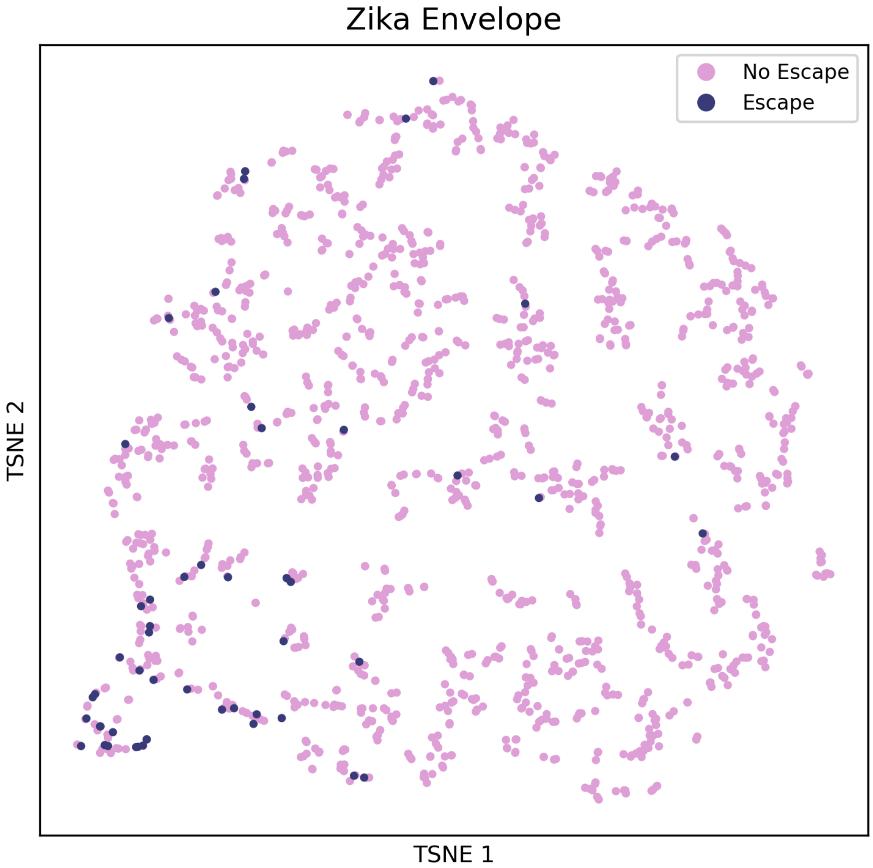


Supplementary Fig. 13: t-SNE of latent embeddings preceding the final linear layer of a multilayer perceptron in the immune escape prediction step for Zika envelope.
